# Beyond Steady-State Adaptation: Evaluating the Dynamic Performance of Biological Feedback Control

**DOI:** 10.64898/2026.09.17.752162

**Authors:** Athena Tamayo-Luisce, Mariana Gómez-Schiavon

**Affiliations:** Laboratorio Internacional de Investigación sobre el Genoma Humano, Universidad Nacional Autónoma de México, Santiago de Querétaro, México; ANID—Millennium Science Initiative Program—Millennium Institute for Integrative Biology (iBio), Santiago 8331150, Chile; Licenciatura en Ciencias Genómicas, Centro de Ciencias Genómicas, Universidad Nacional Autónoma de México, Cuernavaca, México

**Keywords:** Feedback control, Homeostasis, Biochemical adaptation, Transient dynamics

## Abstract

Organisms rely on **feedback control** mechanisms to maintain key biological variables within functional ranges despite persistent perturbations. Understanding how effectively these mechanisms maintain **homeostasis** is fundamental, yet quantitative evaluation of feedback performance remains challenging in nonlinear biological systems. Our previously developed framework, Control Ratio (**CoRa**), addresses this challenge by isolating the contribution of a feedback interaction through controlled comparison with an otherwise identical system in which that interaction has been removed. However, because CoRa evaluates adaptation solely through steady-state responses, it cannot distinguish controllers that ultimately recover to the same state but follow markedly different transient trajectories, despite the potentially profound physiological consequences of those dynamics. Here, we introduce **CoRaDyn**, a framework for evaluating adaptation as a dynamic process rather than solely as a steady-state outcome. Building on the comparative strategy introduced in CoRa, CoRaDyn quantifies the cumulative effect of feedback throughout the post-perturbation response, generating a time-dependent characterization of feedback performance. This approach reveals how the contribution of feedback depends not only on system parameters but also on the time horizon over which adaptation is evaluated, allowing distinct physiological objectives—such as rapid recovery or the generation of transient pulses—to be systematically compared. We demonstrate CoRaDyn using a gene regulatory circuit implementing proportional-integral-derivative (PID) control. Whereas CoRa predicts identical perfect adaptation for all controllers containing integral feedback, CoRaDyn discriminates their transient performance, quantifies the dynamic contributions of proportional and derivative control, and reveals trade-offs that are invisible to steady-state analyses. By extending feedback evaluation from endpoints to the full adaptation process, CoRaDyn broadens the scope of biological questions that can be addressed using the CoRa framework and provides a general approach for comparing feedback architectures when transient dynamics are central to biological function.

**Author summary:** Living cells continuously experience changes in their internal and external environments. Feedback control helps them cope with these disturbances by regulating biological processes to maintain proper function. Two concepts are commonly used to describe these behaviors: **homeostasis**, which emphasizes the ability of a system to recover after a disturbance, and **biochemical adaptation**, which focuses on the transient response that allows cells to detect and respond to environmental changes. Although these phenomena are often studied separately, they can emerge from the same feedback system, simply reflecting different aspects of how feedback control shapes its response to perturbation.

Most existing methods for evaluating biological feedback focus on the final outcome of the response, asking whether a system eventually returns to its original state. However, the path taken to reach that state can be equally important, as different transient responses may have profoundly different physiological consequences. In this work, we present **CoRaDyn**, a computational framework that quantifies the contribution of feedback as the response to a perturbation unfolds over time, rather than only at steady state. Using a gene regulatory circuit with proportional-integral-derivative (PID) control, we show that CoRaDyn distinguishes feedback strategies that appear identical under steady-state analysis while revealing differences in their dynamic performance. By evaluating adaptation as a process instead of only as an endpoint, CoRaDyn provides a general framework for studying how feedback shapes biological behavior across diverse physiological contexts.

## Introduction

Living systems continuously experience changes in both their internal and external environments. To maintain proper function despite persistent perturbations, they rely on **feedback control** mechanisms that regulate key biological variables over time [1–4]. The dynamic behavior of these feedback systems underlies diverse physiological phenomena, making it crucial to understand how feedback control shapes biological behavior.

A major challenge in evaluating biological feedback control is disentangling the contribution of the feedback mechanism itself from the many interacting processes that shape the behavior of nonlinear biological systems. To address this problem, we previously developed the **Control Ratio (CoRa)** framework, which isolates the contribution of feedback by comparing a feedback-controlled system with an otherwise identical system lacking feedback [5]. By quantifying the steady-state response to a perturbation, CoRa provides a rigorous and general framework for evaluating how effectively feedback restores the system after disturbance, independently of the particular nonlinear properties of the underlying biological network. However, because CoRa by design evaluates adaptation through its steady-state outcome, it cannot distinguish systems that achieve the same level of adaptation through fundamentally different transient responses. Hereafter, we use “adaptation” to refer specifically to the dynamical process by which the system approaches its post-perturbation steady state, and “adaptation time” to denote the corresponding timescale. This process can also be viewed as relaxation toward the post-perturbation steady state, but we use “adaptation” here following the terminology of Chevalier et al. [6]. This use of “adaptation” should be distinguished from the broader biological phenomenon of biochemical adaptation, as well as from other uses of the term in different fields.

The importance of considering transient dynamics becomes evident when examining two of the best-known manifestations of biological feedback: homeostasis and biochemical adaptation. **Homeostasis** emphasizes the ability of a system to maintain key biological variables within physiologically relevant ranges despite persistent perturbations (e.g., [7, 8]), whereas **biochemical adaptation** emphasizes the transient response that allows a system to remain responsive to environmental changes while ultimately returning to its pre-perturbation state (e.g., [8–10]). Although these concepts are often discussed independently, both emerge from the same fundamental mechanism: regulating biological variables in changing environments through feedback control. Together, they highlight complementary aspects of the same adaptation process: homeostasis emphasizes the eventual recovery of the system, while biochemical adaptation depends critically on the transient response that precedes it.

This distinction becomes particularly important because, in many biological systems, the transient response is as important as the eventual steady state [11]. Specifically, the speed of adaptation, the transient excursions, or the temporal pattern of the response can shape the resulting physiological outcome [12–16]. As a result, feedback systems that appear equivalent when evaluated solely by their steady-state adaptation may have profoundly different biological consequences. Thus, evaluating biological feedback requires characterizing not only the outcome of adaptation, central to homeostasis, but also the adaptation process itself, which shapes biochemical adaptation.

To address this challenge, we introduce **CoRaDyn**, a framework that builds on the controlled comparison established by CoRa while extending the evaluation of feedback performance from the endpoint of adaptation to the adaptation process itself. Rather than evaluating feedback performance solely through steady-state responses, CoRaDyn quantifies the cumulative contribution of feedback throughout the response to perturbation, generating a time-dependent characterization of feedback performance. This dynamic perspective enables the systematic comparison of feedback systems according to the distinct physiological objectives they must satisfy, including rapid recovery, limited transient deviations, or the generation of transient pulses. We demonstrate the approach using a gene regulatory circuit implementing proportional-integral-derivative (PID) control [6], showing that CoRaDyn distinguishes controllers that exhibit identical steady-state adaptation yet markedly different transient responses, and systematically quantifies the individual contributions of proportional and derivative control to the system’s dynamic performance. By moving beyond steady-state adaptation, CoRaDyn provides a general framework for studying how feedback control shapes biological behavior throughout the adaptation process.

## CoRaDyn Formalism

CoRaDyn builds on the controlled comparison framework established by CoRa [5] to quantify the dynamic contribution of feedback regulation in biological systems. Like CoRa, it relies on mathematical control comparisons [17], evaluating the system of interest against a locally analogous system that is identical in every respect except for the removal of a specific feedback interaction. By isolating the effect of the feedback while preserving the remaining system architecture and operating point, this comparison attributes differences in the response to perturbation exclusively to the feedback mechanism under study.

### Locally Analogous System

To isolate the contribution of a specific feedback interaction to the response dynamics of a biological system, we construct a locally analogous system in which that interaction is removed, hereafter referred to as the feedback-disrupted (*FD*) system. The original and feedback-disrupted systems contain the same biochemical reactions and are described by the same set of differential equations with parameters Θ, differing only in the removal of that interaction (Fig. 1A).

**Fig 1.**
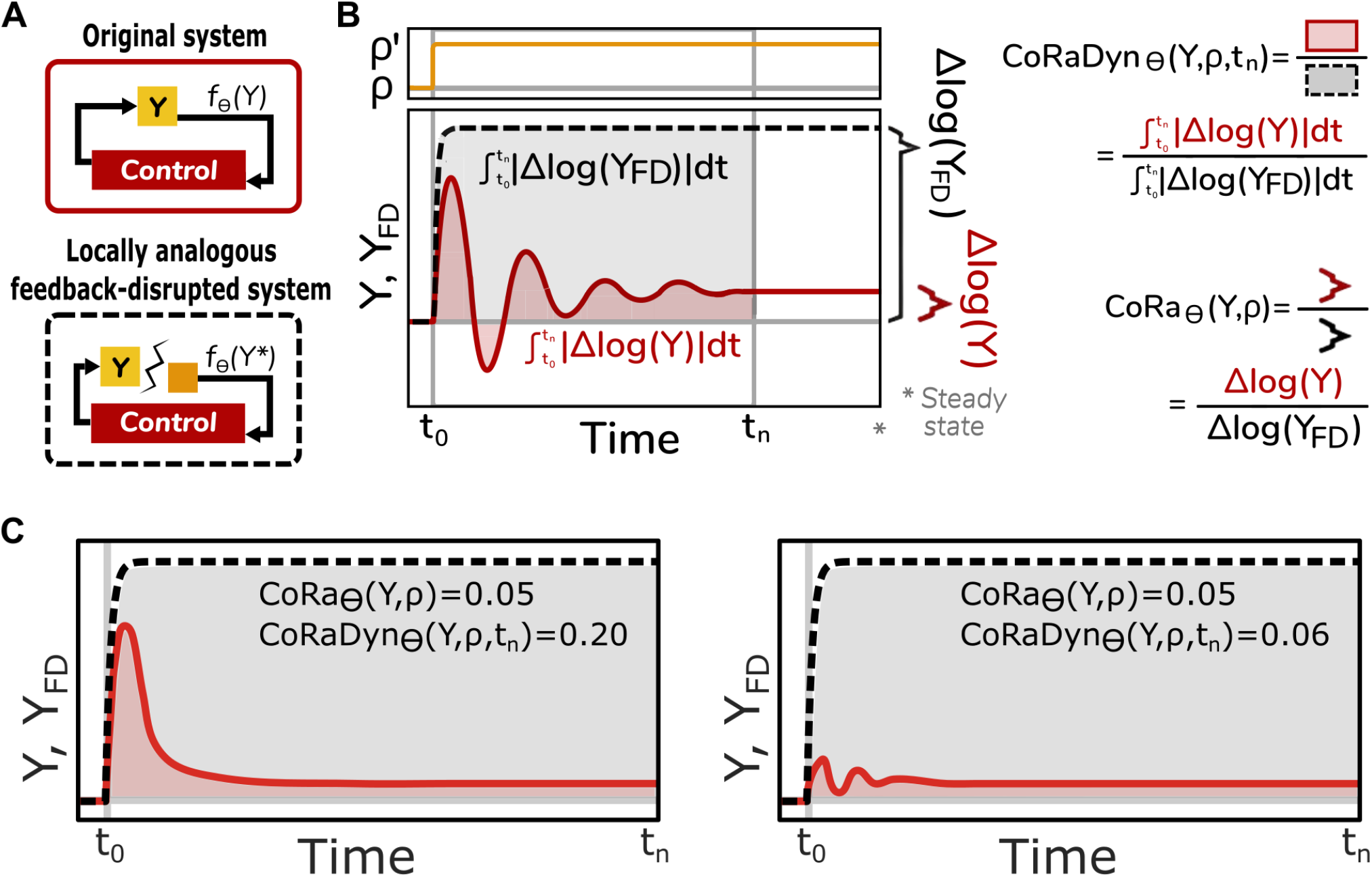
Explaining the CoRaDyn approach. **(A)** Schematic of an original feedback-regulated system (top) and its locally analogous feedback-disrupted counterpart (bottom), constructed by removing the feedback interaction under evaluation while maintaining the same pre-perturbation steady state. The removed interaction is replaced by a constant input chosen so that the feedback signal *f*_Θ_(*Y*) is matched before perturbation. **(B)** Time-course responses of *Y* in the original (red solid line) and feedback-disrupted (black dashed line) systems following a perturbation 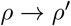. The original CoRa metric quantifies feedback performance from the relative change in steady-state output, CoRa_Θ_(*Y, ρ*) = Δlog(*Y*)*/*Δlog(*Y*_*F D*_). In contrast, CoRaDyn compares the full response trajectories by integrating the absolute logarithmic deviations over time up to an evaluation time *t*_*n*_, 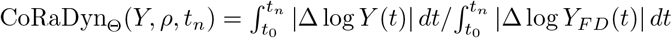. This captures the cumulative contribution of feedback throughout the response to perturbation, rather than only its endpoint. **(C)** Illustration of two systems with identical CoRa values, indicating equivalent feedback performance when assessed solely from steady-state adaptation, but different CoRaDyn values, revealing distinct dynamic contributions of feedback during the adaptation process.

To ensure that the two systems are locally analogous, all reactions and parameter values are preserved (**internal equivalence**), while the removed feedback interaction is replaced by a constant input chosen so that all shared species attain the same pre-perturbation steady state (**external equivalence**). Under these conditions, both systems begin from the same operating point and differ exclusively in the presence or absence of the interaction being evaluated. Consequently, any differences in their responses following a perturbation can be uniquely attributed to the functional contribution of the feedback.

The locally analogous system can be constructed for any individual feedback loop, even in systems containing multiple interacting feedback mechanisms, provided two conditions are satisfied. First, the perturbation must not affect the constant input introduced in the feedback-disrupted system; otherwise, the observed differences cannot be uniquely attributed to the removed feedback interaction. Second, the perturbation must act downstream of the removed feedback interaction and upstream of the controlled species (*Y*), ensuring that the perturbation propagates through the portion of the system regulated by the interaction under study. The construction of locally analogous systems follows the procedure described previously [5], while further details on implementing CoRaDyn are provided in Supplementary Information (*Section S1*).

### From Steady-State to Dynamic Comparison

Up to this point, CoRaDyn follows the same controlled comparison framework as CoRa. The key distinction lies in how the effect of the feedback interaction is quantified. CoRa evaluates feedback performance from the change in the steady-state value of the controlled variable *Y* following a perturbation. In contrast, CoRaDyn quantifies the normalized cumulative effect over the entire response of *Y*, comparing the full response trajectories of the original and feedback-disrupted systems rather than only their final values. This trajectory-level comparison captures differences in the adaptation process, including transient dynamics and temporal delays introduced by the feedback, providing a more comprehensive characterization of feedback performance (Fig. 1B). We denote the parameter being perturbed by *ρ* and the evaluation time by *t*_*n*_, such that CoRaDyn is written as CoRaDyn_Θ_(*Y, ρ, t*_*n*_).

Owing to the normalization inherent to the CoRaDyn formulation (see *Section S1*), its values lie between 0 and 1. As with CoRa, lower values indicate that the feedback attenuates the effect of the perturbation, thereby providing greater control. Specifically, CoRaDyn_Θ_(*Y, ρ, t*_*n*_) ∈ (0, 1) reflects a graded dynamic contribution of the feedback, whereas CoRaDyn_Θ_(*Y, ρ, t*_*n*_) = 1 indicates that the original and feedback-disrupted systems respond identically, implying that the disrupted interaction provides no effective control. Unlike CoRa, however, CoRaDyn cannot attain a value of zero. Because feedback acts only after the perturbation has affected the system, the controlled variable must necessarily undergo a transient deviation from its pre-perturbation steady state. Consequently, the integrated deviation in the numerator of the CoRaDyn expression (Fig. 1B) is always strictly positive. It may, however, become arbitrarily small as the effectiveness of the feedback approaches perfect adaptation and the evaluation time is extended.

A key distinction between CoRa and CoRaDyn is that lower values do not necessarily imply better system performance. Rather, the interpretation of CoRaDyn depends on the regulatory objective of the system under study. Systems optimized to minimize deviations from their operating point generally exhibit lower CoRaDyn values, whereas systems that require a transient response before returning to equilibrium may exhibit larger values at early evaluation times that decrease as adaptation proceeds. Thus, unlike CoRa, which evaluates only the endpoint of adaptation, CoRaDyn captures differences in the dynamic response to perturbation and can distinguish regulatory strategies with similar steady-state outcomes (Fig. 1C).

### Time-Dependent Characterization of Feedback Performance

The evaluation time *t*_*n*_ defines the temporal window over which the dynamic contribution of feedback is quantified. Although CoRaDyn can be evaluated at a single time point, its full temporal behavior is revealed by computing the metric across a range of evaluation times. This generates a time-dependent profile of feedback performance, CoRaDyn_Θ_(*Y, ρ, t*), which captures how the feedback contribution evolves throughout the system’s perturbation response (Fig. 2A).

**Fig 2.**
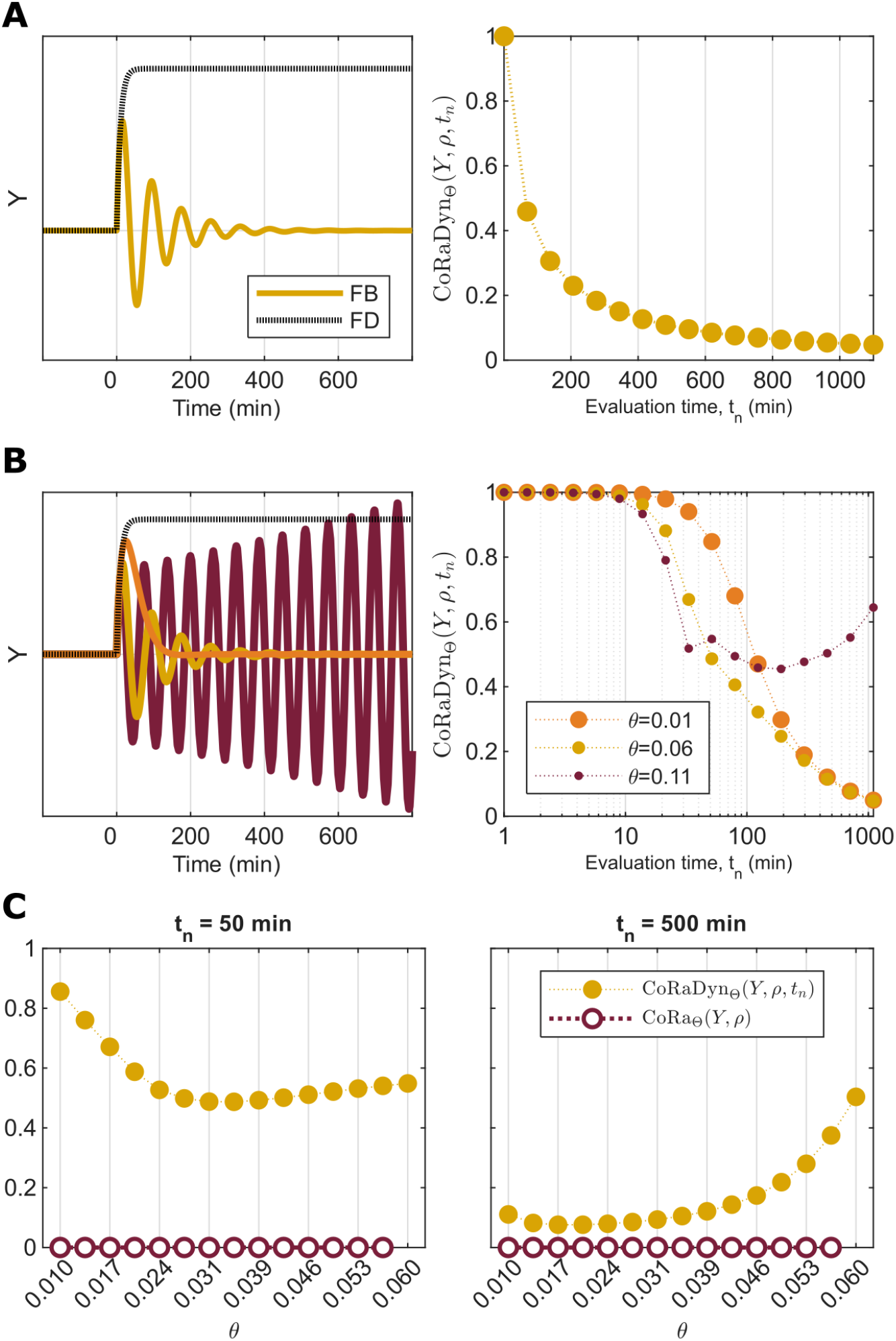
Time-dependent characterization of feedback performance using CoRaDyn. **A:** Example of a CoRaDyn temporal profile, obtained by evaluating feedback performance over increasing evaluation times. On the left, the dynamic responses of the original (solid red) and feedback-disrupted (dashed black) systems are shown; on the right, the corresponding change in CoRaDyn as the evaluation time (*t*_*n*_) increases is shown. This example corresponds to a system with perfect adaptation, for which the steady-state CoRa value is zero. **B:** Example of how the CoRaDyn temporal profile varies as a parameter *θ* ∈ Θ changes. On the left, the dynamic responses of the original (solid lines) and feedback-disrupted (dashed lines) systems are shown for three values of *θ*; on the right, the corresponding temporal profiles of CoRaDyn are shown. **C:** Examples of CoRaDyn values evaluated at short (*t*_*n*_ = 50 *min*; left) and long (*t*_*n*_ = 500 *min*; right) evaluation times, together with the corresponding CoRa values, as a parameter *θ* varies. For the largest value of *θ* shown, the system exhibits sustained oscillations; therefore, CoRa cannot be defined, whereas CoRaDyn remains applicable. See *Section S2* for equations and Table S2 for parameter values.

At times immediately following the perturbation, the original and feedback-disrupted systems remain similar due to their locally analogous construction, resulting in values of CoRaDyn close to one. As the feedback interaction begins to influence the system dynamics, differences between the two trajectories emerge, and CoRaDyn decreases according to the extent and timescale of the corrective response. Thus, the temporal profile of CoRaDyn provides information about how feedback contributes over time, rather than only whether the system eventually reaches a controlled state.

As evaluation time increases, CoRaDyn approaches the steady-state contribution of feedback captured by CoRa. For systems that reach a new steady state following the perturbation, increasing *t*_*n*_ progressively reduces the relative contribution of transient differences and causes CoRaDyn to converge toward the corresponding CoRa value. In contrast, for systems exhibiting persistent oscillations or other non-steady-state behaviors, this convergence is not expected, and the temporal profile itself becomes the relevant characterization of feedback performance.

Beyond time-dependent analyses, CoRaDyn can also be systematically evaluated across parameter ranges to visualize how feedback performance changes with the underlying system properties. For a parameter *θ* ∈ Θ, computing CoRaDyn_Θ_(*Y, ρ, t*_*n*_) across different values of *θ* generates a dynamic characterization of feedback performance across parameter space (Fig. 2B-C). For each parameter value, the feedback-disrupted system must be reconstructed to preserve the conditions of a mathematically controlled comparison. Together, parameter- and time-dependent analyses provide a multidimensional view of feedback function, revealing how regulatory architectures shape both the magnitude and timescale of control.

Finally, unlike CoRa, which is defined in terms of local steady-state changes and is therefore insensitive to perturbation magnitude in the limit of sufficiently small perturbations, CoRaDyn may exhibit some dependence on Δ*ρ*. For most systems and sufficiently small perturbations, this dependence is minimal. However, because CoRaDyn captures transient dynamics, its value may vary when perturbations alter the qualitative response trajectory, particularly near bifurcation points. For example, perturbations of different magnitudes may induce qualitatively distinct behaviors, such as damped versus sustained oscillations, resulting in different temporal profiles and CoRaDyn values (Fig. S1). Therefore, while maintaining the perturbative regime required for controlled comparison, perturbation magnitude should be considered when interpreting dynamic feedback performance, particularly in systems operating near dynamical transitions.

### Stress-testing CoRaDyn with biomolecular PID control

As a stress-test for the application and power of CoRaDyn, we consider the proportional-integral-derivative (PID) controller implemented with biomolecular components, originally proposed by Chevalier et al. [6] (Fig. 3A). PID control provides an ideal benchmark because it combines feedback mechanisms with distinct and potentially competing dynamical objectives: integral feedback ensures perfect adaptation, whereas proportional and derivative feedback regulate the speed and stability of the response. Thus, although all PID configurations can potentially achieve the same adapted steady state, their transient behaviors can differ substantially—differences that are invisible to endpoint measures such as CoRa, but can be captured by CoRaDyn.

**Fig 3.**
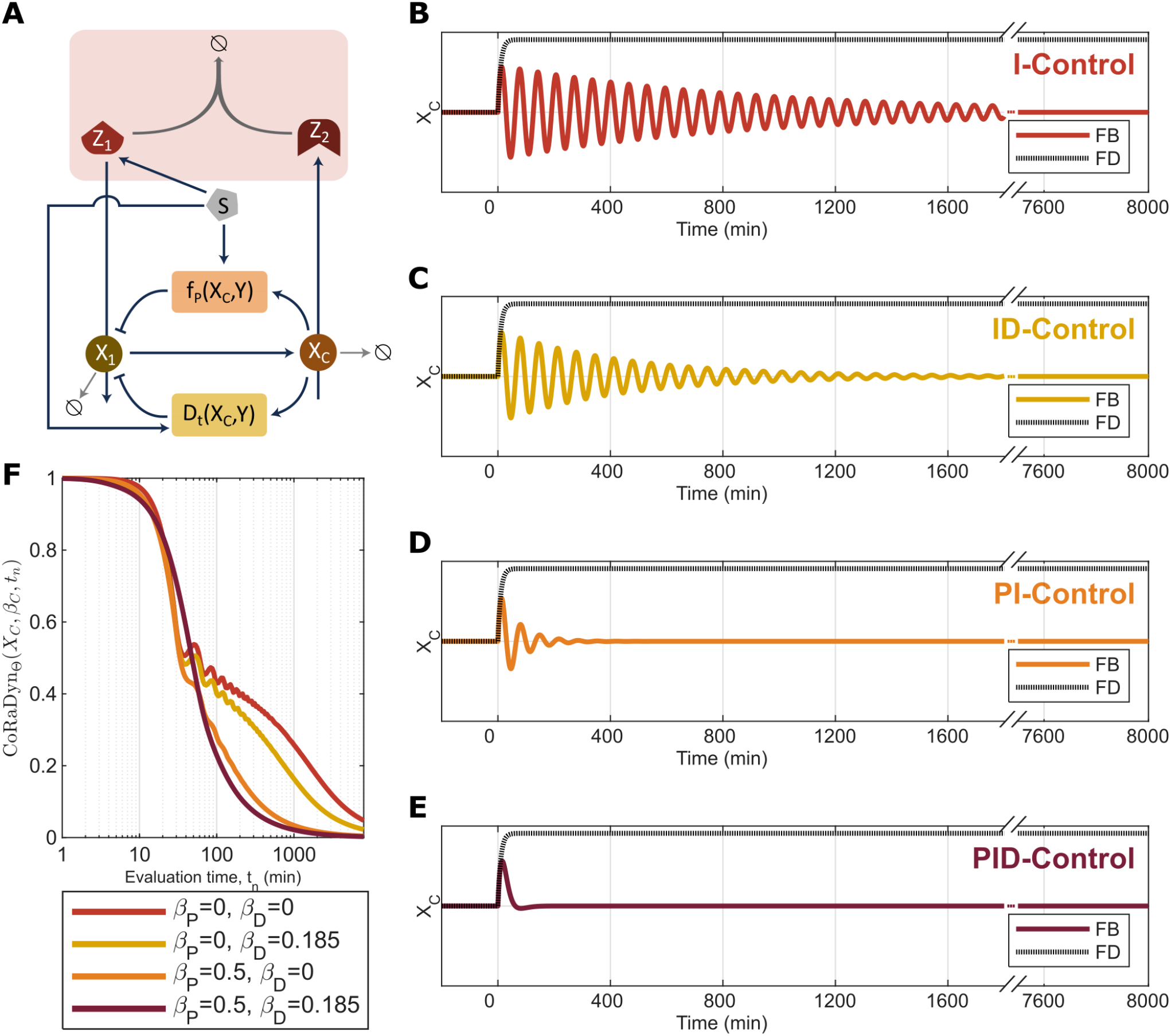
CoRaDyn reveals differences in transient dynamics beyond steady-state adaptation. **A:** Schematic representation of the biomolecular PID controller analyzed in this study (adapted from Chevalier et al., 2019). The integral (I), proportional (P), and derivative (D) feedback modules are highlighted in red, orange, and yellow, respectively. **B–E:** Dynamic responses to a step perturbation for four controller configurations: (**B**) integral-only (I; *β*_*P*_ = 0, *β*_*D*_ = 0), (**C**) integral-derivative (ID; *β*_*P*_ = 0), (**D**) integral-proportional (PI; *β*_*D*_ = 0), and (**E**) full PID control. All configurations exhibit perfect adaptation and therefore have identical steady-state CoRa values (CoRa = 0), despite their distinct transient responses. **F:** CoRaDyn as a function of the evaluation time (*t*_*n*_) for the four controller configurations shown in panels B–E. In all cases, CoRaDyn converges to zero as the systems approach their adapted steady states, but the trajectories differ substantially during the transient response. Differences in the initial decay of CoRaDyn and changes in the relative ranking of controller performance over time are observed. See *Section S2* for equations and Table S2 for parameter values.

The integral component of the controller is implemented through the antithetic feedback (ATF) motif introduced by Briat et al. [18]. In this architecture, two molecular species mutually annihilate through binding, generating an integral control action that drives the steady-state error to zero under appropriate conditions. Consequently, the ATF motif provides the integral control action required for perfect adaptation after a step perturbation (i.e., a CoRa of zero). However, depending on the parametrization, the system may converge to the adapted steady state through slow transients, or, in some cases, may fail to settle altogether and exhibit sustained oscillations. Thus, achieving perfect adaptation and satisfying physiological performance objectives are distinct properties of the controller.

The proportional and derivative components address these limitations by shaping the transient response. Proportional feedback responds to the magnitude of the deviation from the target state, accelerating recovery, whereas derivative feedback responds to the rate of change of this deviation, reducing overshoot and improving stability [19]. Therefore, the relative contribution of each feedback component depends on the temporal scale at which performance is evaluated—a feature that motivates the dynamic assessment provided by CoRaDyn.

In the model considered here, each control strategy is governed by a tunable weight—*β*_*I*_ (integral), *β*_*P*_ (proportional), and *β*_*D*_ (derivative)—that determines its contribution to the overall controller. Moreover, each feedback loop can be selectively disrupted, enabling the controlled comparison of locally analogous systems and the quantification of individual feedback contributions using the CoRaDyn framework (see *Section S2* for implementation details). In the following sections, we use the PID controller model of Chevalier et al. [6] to illustrate three key capabilities of CoRaDyn: evaluating feedback performance across time, mapping the dependence of adaptation performance on control parameters, and dissecting the contribution of individual feedback components within a composite regulatory architecture.

#### Feedback contribution depends on the temporal scale of evaluation

To illustrate the importance of evaluation time, we compared four configurations of the biomolecular PID controller that achieve perfect adaptation: integral-only control (I; *β*_*P*_ = 0, *β*_*D*_ = 0), integral-derivative control (ID; *β*_*P*_ = 0), integral-proportional control (PI; *β*_*D*_ = 0), and the full PID controller. The proportional-derivative (PD) configuration is not included because, in the absence of integral control, it does not achieve perfect adaptation. For each configuration, we simulated the response to an identical perturbation and computed CoRaDyn as a function of the evaluation time, *t*_*n*_ (Fig. 3).

Because all four architectures contain the same integral control module, they achieve perfect adaptation following the perturbation and therefore have identical steady-state performance (CoRa = 0). Consequently, CoRa cannot distinguish among these controllers despite their markedly different transient responses (Fig. 3B–E). CoRaDyn, in contrast, quantifies feedback performance throughout the adaptation process, revealing differences that are invisible to endpoint-based measures.

The resulting CoRaDyn trajectories (Fig. 3F) reveal distinct temporal profiles of feedback performance for the four controller architectures. In all cases, CoRaDyn decreases toward zero as the evaluation time increases, reflecting the progressive recovery of the perturbed system toward its adapted steady state. However, the rate of this decrease differs substantially across controllers, demonstrating that proportional and derivative feedback alter the transient contribution of feedback even though they do not affect the final adapted state, consistent with the results of Chevalier et al. [6]. At early evaluation times, the ID controller exhibits the lowest CoRaDyn values, consistent with its ability to reduce the initial transient response. During the same period, the I and PI controllers display similar trajectories. As the response progresses, however, the PI controller surpasses the I controller, eventually catching up with the ID controller and reaching lower CoRaDyn values more rapidly. The full PID controller ultimately outperforms all other configurations, although this advantage only becomes apparent after the initial phase of the response.

Importantly, the relative performance of the different controllers is not constant over time. The CoRaDyn trajectories are non-monotonic with respect to one another, such that the controller exhibiting the strongest feedback contribution depends on the evaluation time selected. Thus, no single architecture can be considered uniformly superior without specifying the temporal objective of interest. This result highlights a key advantage of CoRaDyn. Because it explicitly incorporates the evaluation time, feedback architectures can be compared according to the timescale over which adaptation is required, rather than solely according to their eventual steady-state behavior.

#### Evaluation time reveals distinct roles for proportional, integral, and derivative feedback

The examples presented in Fig. 3 illustrate that the apparent contribution of proportional and derivative control depends strongly on the evaluation time. However, these conclusions correspond to a single parameterization of the PID controller and therefore should not be interpreted as general properties of the controller architecture. A key advantage of the CoRaDyn framework is that it enables systematic exploration of parameter space, allowing feedback performance to be compared across controller configurations and temporal objectives.

To verify that CoRaDyn remains consistent with established measures of transient performance, we first evaluated CoRaDyn at a fixed evaluation time (*t*_*n*_ = 500 min) across the PID parameter space. The resulting performance landscapes closely resemble those previously obtained using adaptation time [6] (Fig. S2). This agreement confirms that, when restricted to a single evaluation time, CoRaDyn captures the same qualitative trends identified using conventional transient metrics.

Unlike adaptation time, however, CoRaDyn is not limited to a single temporal summary of the response. Because it quantifies cumulative feedback performance over an arbitrary evaluation interval, the same collection of simulations can be interrogated across multiple temporal scales. Furthermore, whereas adaptation time becomes undefined when the perturbed system fails to converge within the observation window, CoRaDyn remains applicable to sustained post-perturbation oscillations. The parameter combinations for which CoRaDyn cannot be evaluated (white/empty regions in Fig. S2) instead correspond to systems lacking a stable pre-perturbation steady state. As with adaptation time and CoRa, a stable pre-perturbation steady state is required to define the reference condition (see Table S1).

To systematically investigate how the contribution of each control strategy changes over time, we summarized CoRaDyn across the explored controller parameter space using the median over the remaining controller weights. We used the median as a robust summary of these distributions, which can span a broad range of CoRaDyn values (the complete distributions are shown as boxplots in Figs. S3–S14). Specifically, we computed 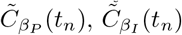, and 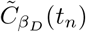, corresponding to the median CoRaDyn obtained for fixed proportional (*β*_*P*_), integral (*β*_*I*_), and derivative (*β*_*D*_) control weights, respectively, while taking the median over all combinations of the remaining two controller weights (Fig. 4A; see caption for formal definitions). In all cases, the median CoRaDyn decreases with increasing evaluation time, reflecting the progressive recovery of the perturbed systems. However, the dependence of CoRaDyn on the individual controller weights varies substantially throughout the temporal profile.

**Fig 4.**
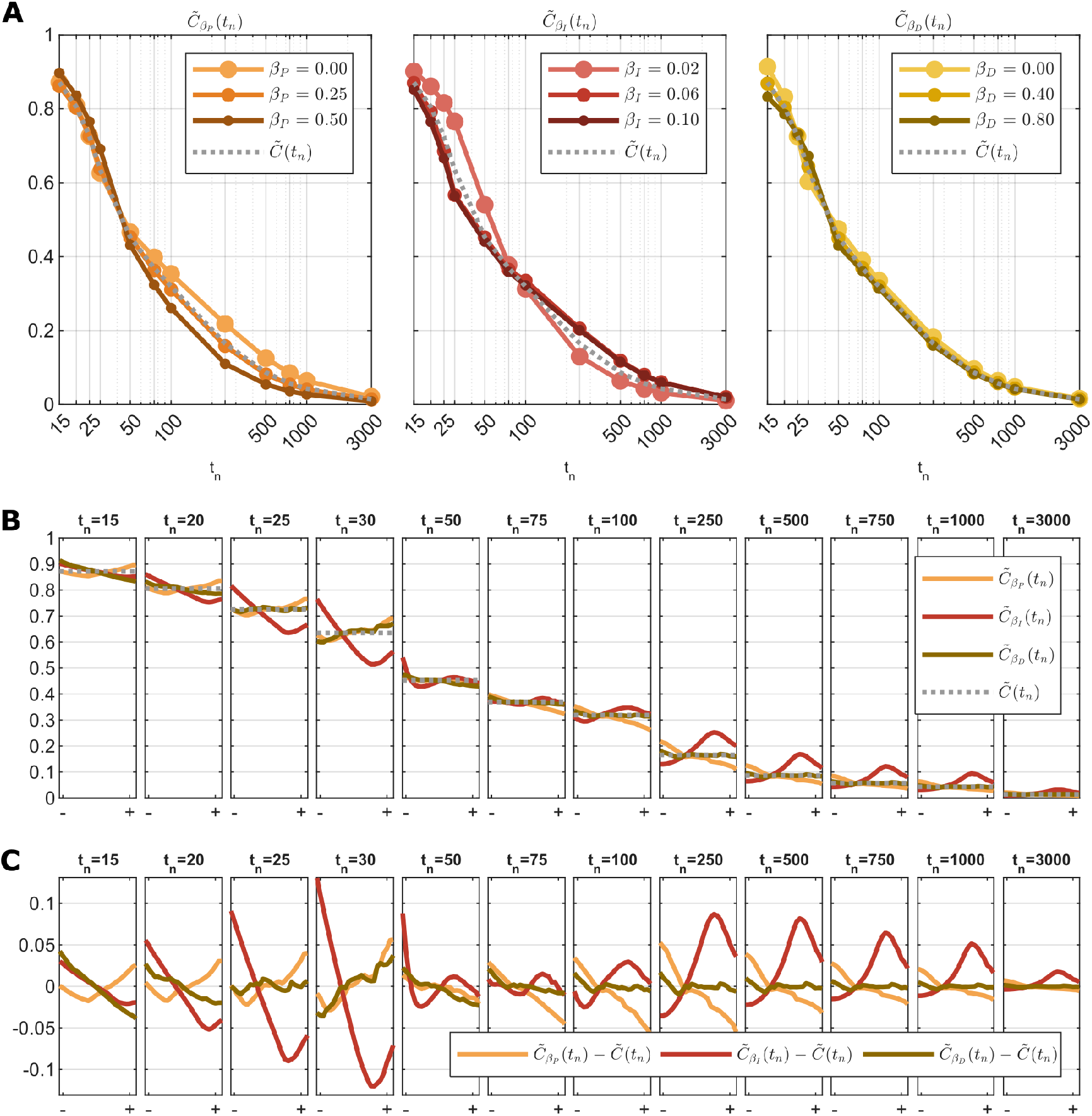
Systematic evaluation of the PID controller across parameter space reveals distinct qualitative effects at different evaluation times. **A:** Temporal profiles of the median CoRaDyn as a function of the evaluation time (*t*_*n*_; logarithmic scale). Each colored curve corresponds to the median CoRaDyn over all combinations of the remaining two controller weights for a fixed value of the indicated control weight, i.e., 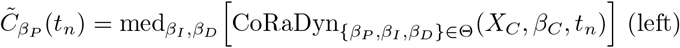, 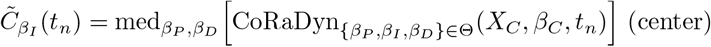, and 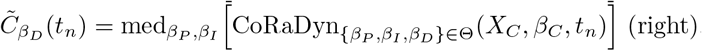. The dotted gray line denotes the global median across all controller weight combinations, 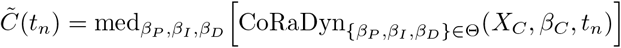. **B:** Median CoRaDyn as a function of each controller weight for representative evaluation times (indicated above each panel). Curves show 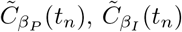, and 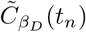 as the corresponding controller weight varies, from low (−) to high (+) values within the explored parameter range, while the dotted gray line indicates the global median 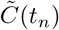 for that evaluation time. **C:** Same data as in **B**, shown as deviations from the global median. Curves represent 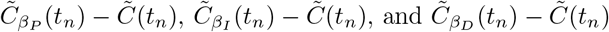, highlighting the relative contribution of each control component with respect to the overall median CoRaDyn at each evaluation time. Fig. S15 presents the same data reorganized by controller weight to facilitate comparison across evaluation times. See *Section S2* for equations and Table S2 for parameter values.

At early evaluation times (*t*_*n*_ = 15 min), increasing the derivative control weight consistently decreases 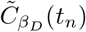, indicating that derivative control contributes most strongly during the initial stages of the response. In contrast, the dependence on the proportional control weight is markedly non-monotonic, with intermediate values of *β*_*P*_ yielding the lowest values of 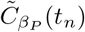. Similar non-monotonic relationships emerge at many other evaluation times and for all three controller weights, demonstrating that the dynamic contribution of each feedback component cannot be characterized by a simple monotonic dependence on its strength.

The influence of the integral controller becomes particularly evident at longer evaluation times. For example, at *t*_*n*_ = 250 min, the smallest values of 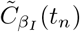 occur for the lowest values of *β*_*I*_, whereas larger—but not maximal—integral weights produce the largest values of 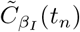. Moreover, *β*_*I*_ often generates the widest spread in 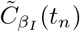 among the three controller weights (Fig. 4C), indicating that the contribution of integral feedback becomes increasingly sensitive to parameter selection as longer timescales are considered.

Together, these analyses demonstrate that the apparent importance of the proportional, integral, and derivative feedback components depends jointly on their control weights and on the temporal objective used to evaluate performance. Rather than providing a single measure of adaptation quality, CoRaDyn enables systematic exploration of how distinct controller architectures satisfy different dynamic objectives, revealing trade-offs that remain inaccessible to endpoint-based metrics.

#### Dissecting the Contribution of Individual Feedback Loops

Finally, we demonstrate how CoRaDyn inherits one of the central principles of the original CoRa framework: the use of locally analogous systems to isolate the contribution of individual feedback loops while preserving the remainder of the network unchanged. Although this possibility was discussed in the original formulation of CoRa, it was not explicitly demonstrated. Here, we extend this principle to dynamic feedback evaluation, allowing the contribution of multiple feedback mechanisms to be quantified within a single controller architecture.

To illustrate this capability, we consider a fully configured PID controller and construct four locally analogous systems by selectively disrupting (1) proportional feedback, (2) derivative feedback, (3) both proportional and derivative feedback, or (4) all feedback loops, and apply CoRaDyn to each resulting comparison (Fig. 5). In contrast to the comparisons in Fig. 3, which contrasted distinct controller architectures, all controlled comparisons here are performed on the same underlying system and parameterization. Consequently, differences in CoRaDyn can be directly attributed to the specific feedback loop(s) disrupted.

**Fig 5.**
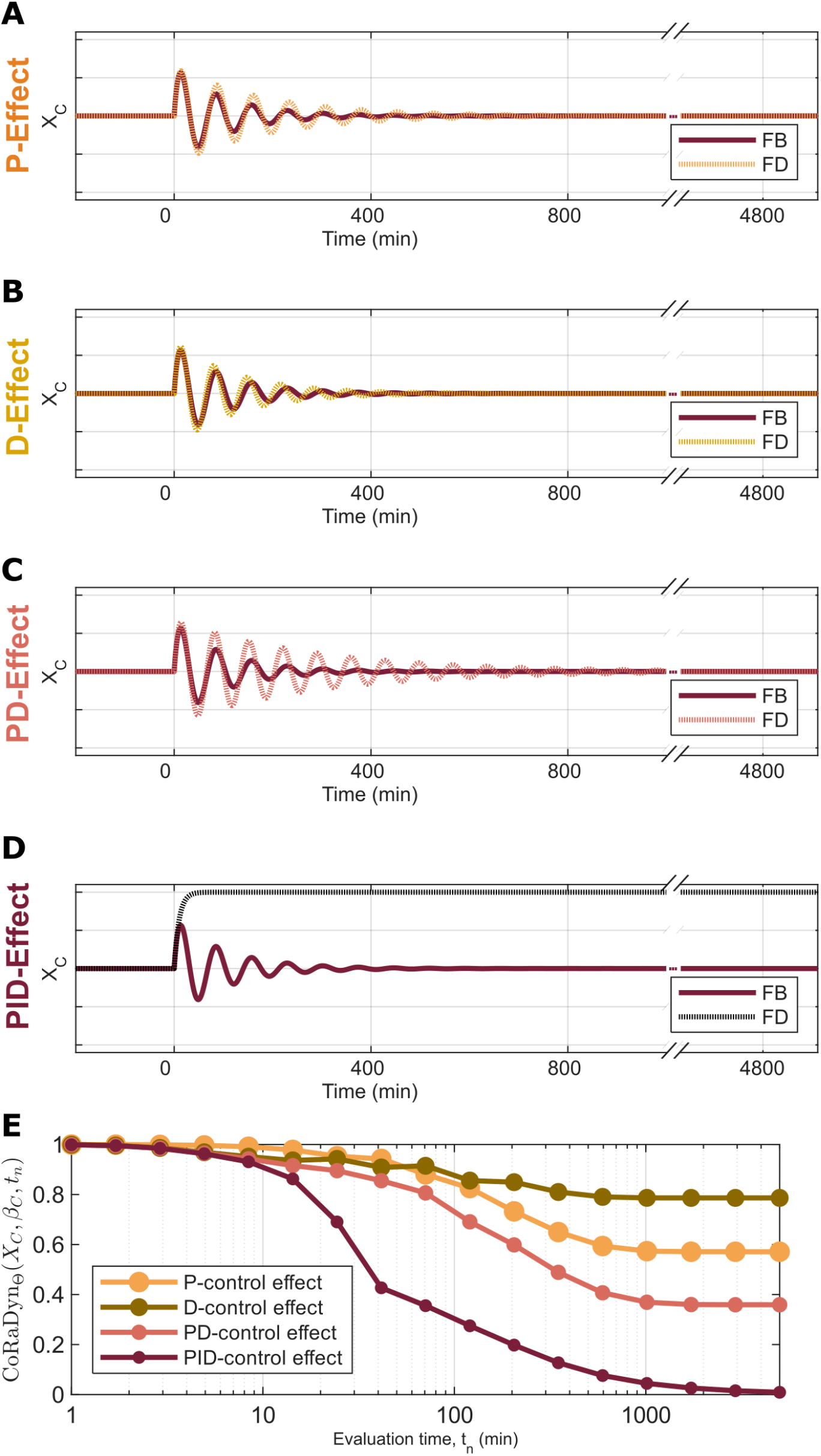
CoRaDyn isolates the dynamic contribution of overlapping feedback loops. **A–D:** Dynamic response of the controlled variable (*X*_*C*_) following a step perturbation over *β*_*C*_ ∈ Θ applied at time zero. In each panel, the original PID controller (dark red) is compared with its corresponding locally analogous system, in which selected feedback loops have been disrupted while preserving the remaining network unchanged: (**A**) proportional feedback removed (orange), (**B**) derivative feedback removed (yellow), (**C**) proportional and derivative feedback removed (red), and (**D**) all feedback loops removed (black). **E:** CoRaDyn as a function of the evaluation time (*t*_*n*_) for the controlled comparisons shown in panels A–D, using the same color scheme. Because the locally analogous systems in panels A–C retain integral control, they exhibit perfect adaptation despite their altered transient responses. Consequently, CoRaDyn quantifies the cumulative dynamic contribution of the disrupted feedback loop(s), enabling the individual and combined effects of proportional, derivative, and integral control to be compared within the same controller architecture. See *Section S2* for equations and Table S2 for parameter values. An alternative parametrization can be observed in Fig. S16.

For each controlled comparison, we computed CoRaDyn as a function of the evaluation time (Fig. 5E). Because the locally analogous systems in which only proportional and/or derivative feedback is disrupted retain integral control, they still exhibit perfect adaptation despite their altered transient responses. Consequently, the observed differences do not reflect changes in steady-state adaptation, but rather differences in the transient correction of the perturbation. Since CoRaDyn is evaluated relative to the corresponding locally analogous system, each trajectory quantifies the fraction of the perturbation remaining in the intact controller relative to the response expected in the absence of the disrupted feedback loop. Thus, lower CoRaDyn values indicate that the removed feedback mechanism contributed more strongly to correcting the perturbation. In contrast, the comparison in which all feedback loops are disrupted lacks integral control and therefore exhibits a sustained deviation from the adapted steady state.

The resulting CoRaDyn profiles reveal distinct contributions of the three control strategies. For the parameterization considered here, removing proportional feedback produces lower CoRaDyn values than removing derivative feedback for most evaluation times, indicating a greater contribution of proportional control to overall dynamic performance, although derivative control has a greater contribution during the earliest phase of the response. The CoRaDyn profiles associated with proportional and derivative feedback intersect repeatedly, demonstrating that their relative contributions change throughout the transient response. Thus, although proportional feedback dominates the cumulative response in this example, derivative feedback provides the greater benefit over specific temporal windows (Fig. 5E).

As in the previous analyses, these conclusions depend on the parameterization of the controller, and an additional example is provided in Fig. S16. Nevertheless, the controlled comparison framework enables the contribution of each feedback mechanism to be interpreted directly from the CoRaDyn trajectories, providing a principled decomposition of dynamic feedback performance that is inaccessible to steady-state analyses alone. In particular, whereas CoRa assigns identical values to all controllers exhibiting perfect adaptation, CoRaDyn reveals how individual feedback loops contribute to different phases of the adaptive response, thereby extending the controlled comparison principle of CoRa from steady-state adaptation to transient dynamics.

## Discussion

### CoRaDyn extends the controlled-comparison approach of CoRa beyond steady-state adaptation to full response trajectories

The Control Ratio (CoRa) framework was designed to systematically evaluate feedback control in modeled biological systems [5], comparing the steady-state of the original system with that of a locally analogous system in which the feedback under study is disrupted. Its focus on steady-state adaptation, however, means that any two controllers sharing the same proportional relationship between their feedback and feedback-disrupted steady-state deviations will be assigned the same CoRa value, regardless of the timescale or shape of their underlying transient dynamics (Fig. 3B–E).

Such differences can be biologically consequential. While some systems need to ensure **homeostasis**, minimizing deviations from a physiologically appropriate state when disturbed [1–3], others must instead achieve physiological function through their dynamics, such as a particular response speed [16], a specific dynamical pattern [12–14], or the generation of transient excursions [15], properties most associated with **biochemical adaptation**. Although homeostasis and biochemical adaptation are both manifestations of feedback control, a metric that relies only on steady-state endpoints is less suited to characterize the latter.

CoRaDyn overcomes these limitations by extending CoRa’s controlled-comparison framework to assess a system’s cumulative perturbation response over a specified evaluation time (*t*_*n*_), rather than a single steady-state endpoint. Taking the transient response into account allows differences in the performance of controllers that are indistinguishable to CoRa to be quantified (Fig. 2C). Conducting the assessment at different evaluation times further reveals how the relative performance of controllers that differ in parametrization or structure shifts as the evaluation window varies (Fig. 3F, Fig. 5E). Most importantly, it also allows the individual contributions of distinct feedback components to be isolated over time (Fig. 5E).

### Feedback contributions depend on the temporal scale of evaluation

The contribution of a given feedback interaction to a system’s perturbation response varies with the evaluation time (*t*_*n*_) (Fig. 2A), and different feedback mechanisms can occupy distinct temporal roles within the same response (Fig. 5E). When CoRaDyn temporal profiles from controllers with different parametrizations or structures are overlaid, the times at which they cross, converge, or separate mark specific phases of the response during which particular feedback components contribute the most. The overall shape of a profile further indicates whether a given contribution is sustained, transient, or changes character over the course of adaptation (Figs. 3-5) — information that is lost whenever the evaluation time is fixed rather than explored.

Furthermore, CoRaDyn temporal profiles also reveal that relative performance can change non-monotonically across the parameter space (Fig. 4B-C). This means that a given set of parameters can strengthen a feedback’s contribution at early evaluation times while weakening it over longer periods, or the reverse. One should therefore be cautious of interpreting the effect of a feedback mechanism and associated parameters as globally beneficial or detrimental, and instead interpret it within the context of the specific temporal scale and parameter selection over which feedback control is being evaluated.

Additionally, this temporal perspective enables the evaluation of feedback control in systems that never settle into a post-perturbation steady state (Fig. 2; Fig.S16C). Such cases are worth analyzing since biological oscillators may be capable of showing robust homeostatic and adaptive behaviors (Thorsen et al., 2014). In the presence of persistent oscillations, however, a steady-state measure such as CoRa cannot be implemented, because no steady state is reached. By not requiring convergence to a post-perturbation steady state, CoRaDyn instead allows post-perturbation persistent oscillations to be characterized.

### CoRaDyn enables systematic comparisons across parameter conditions and isolates the contributions of individual feedback components

The molecular PID system proposed by Chevalier et al. [6] illustrates why considering the full perturbation response trajectory, rather than a single steady-state endpoint, matters when evaluating feedback control. The system consists of proportional (P), integral (I), and derivative (D) feedback modules whose relative contributions can be tuned through individual weights. Given that every parameter combination explored here drives the system to the same steady state or to sustained oscillations after a step perturbation, CoRa cannot distinguish between parameter sets that produce different transient responses. This is because it returns zero for every set of parameters that leads to adaptation, and becomes undefined whenever the system oscillates instead. CoRaDyn, in contrast, detects differences in the perturbation response as different weights are assigned to the three modules (Fig. 3F). Because the effect of a given module, however, depends jointly on its own parameters and on those of the other modules, any single example like this one is highly dependent on the particular conditions evaluated, and extracting general trends instead requires a systematic exploration of the parameter space (Fig. 4).

Such systematic analysis is allowed by CoRaDyn, and the resulting trends are broadly consistent with what is known about the PID system studied by Chevalier et al. [6], while also providing additional nuance (Fig. 4; Fig. S2-S15). Proportional control is known to increase the speed of the response, and accordingly, high proportional weights lower CoRaDyn values once the evaluation window is long enough to capture the reduced settling time. However, early in the response very high weights also increase CoRaDyn values, consistent with an increase in overshoot. The integral weight’s median effect also reverses as the evaluation time increases. Strong integral action corrects deviations quickly at the outset, lowering CoRaDyn early on, but the same action can also enlarge the settling time or promote oscillatory behavior, thereby raising CoRaDyn once the evaluation window is long enough to capture this transition. Derivative control, on the other hand, is known to reduce overshoot and speed convergence to steady state, consistent with its dominant contribution during the initial stages of the response. However, only low weights continue to contribute non-negligibly to perturbation attenuation as time progresses, suggesting that its contribution to convergence speed is contextual. Together, while the trends observed for proportional and derivative control align with previous descriptions in the literature, none of the three control weights, proportional, integral, or derivative, improve the response monotonically as they increase. Each instead has evaluation-time windows and parameter ranges over which its median contribution may be considered to be positive, negative, negligible, or to entail trade-offs.

Beyond capturing how parameter variation reshapes the system’s behavior, CoRaDyn allows quantifying the individual contributions of proportional and derivative control over time (Fig. 5E) — a capability proposed in the original CoRa paper but never demonstrated [5]. By selectively disrupting the proportional and derivative feedback, both jointly and individually, CoRaDyn shows that each can have qualitatively distinct, time-dependent effects on the response. Interestingly, the temporal profiles of the systems individually lacking proportional or derivative feedback intersect, indicating that their relative contributions reverse over time, with derivative control contributing more at the response onset and proportional control contributing more at longer times. These temporal profiles, however, depend on the particular parametrization, as illustrated by the different patterns observed under alternative parametrizations (Fig. S16). Thus, the way in which proportional and derivative feedback shape the response depends jointly on the temporal context and the parameters of the system.

### Broader implications and future directions

CoRaDyn broadens the class of biological questions that can be addressed through the mathematically controlled comparison of feedback systems. Its temporal profiles identify when feedback contributes to adaptation, how that contribution changes over the course of the response, and how the relative advantage of different feedback mechanisms shifts depending on the evaluation time considered. This perspective may be useful for systems in which transient dynamics are functionally important, including, for example, regulatory processes in which the timing or amplitude of a response shapes downstream physiological outcomes.

An important consequence of these results is that no single feedback architecture is likely to be optimal independently of the temporal objective under consideration. Different mechanisms contribute preferentially at different stages of the response, and the non-monotonic dependence of CoRaDyn on evaluation time shows that the apparent advantage of a feedback strategy can shift as the adaptation process unfolds. CoRaDyn makes these temporal trade-offs explicit rather than obscuring them in a single summary value.

As with CoRa, the interpretation of a CoRaDyn value ultimately depends on the system being interrogated, its biological function, and the question motivating the analysis. A regulatory circuit that maintains distinct pH levels across the cytosol and other cellular compartments [3], for example, may exhibit low CoRaDyn values throughout the response, since minimizing deviations from its operating point is itself the relevant function. A circuit that regulates bacterial chemotaxis [15], in contrast, may exhibit high CoRaDyn values when evaluated at intermediate times.

However, this does not signify inappropriate performance, but rather reflects a function that depends on a transient response with a substantial initial deviation. This context dependence is not a shortcoming of the metric but a structural feature of it, requiring the user to state explicitly what a well-adapted response should look like for the biological system under study.

The choice of evaluation time should likewise be treated as part of the definition of the control objective rather than as an arbitrary modeling choice. Candidates for *t*_*n*_ include the time by which the perturbing signal is expected to resolve or the timescale over which the biological process of interest normally unfolds (e.g., the duration of a relevant developmental or signaling window, or the timescale over which experimental measurements are available). However, as shown, inspecting the full CoRaDyn temporal profile rather than committing to a single fixed evaluation time in advance may be more informative.

Several extensions of this framework could further broaden its applicability. Incorporating stochastic dynamics would allow feedback contributions to be evaluated in systems where intrinsic and extrinsic fluctuations are central to the response. CoRaDyn could also be applied to more complex regulatory architectures, used to compare regulatory strategies across less theoretical biological systems, and applied to the systematic design or optimization of feedback strategies under explicitly defined temporal objectives. Additionally, although CoRaDyn does not require a post-perturbation steady state, it still depends on a pre-perturbation one, so it cannot currently be applied to biological oscillators that achieve homeostatic or adaptive behavior around a periodic rather than a stationary reference state [20]. Thus, extending the locally analogous comparison framework to accommodate a periodic pre-perturbation state represents a further direction for future work.

## Conclusion

In summary, by comparing the full response trajectories of two otherwise identical systems that differ only in the presence of a given feedback interaction, rather than only relying on their final steady states, CoRaDyn captures information about feedback performance that CoRa does not consider by construction. This shift makes it possible to ask not only whether a given feedback matters for a system’s return to its pre-perturbation steady state after a disturbance, but when it matters throughout the response, and whether trade-offs arise along the way. Accordingly, applied to a molecular PID controller, CoRaDyn was able to extract the individual effects of different feedback components on the system’s transient behavior, capturing when each component contributes, how that contribution evolves over time, and where its effects trade off against those of the others. These are relevant capacities for evaluating systems where the path they take to recover from a perturbation is as physiologically consequential as the steady-state endpoint itself.

Beyond characterizing existing circuits, this temporal framework has a natural application in their design. As synthetic biology seeks specific transient behaviors — such as a bounded overshoot, a fast settling time, or a sustained pulse — rather than steady-state robustness alone, CoRaDyn offers a way to score candidate controller architectures against these temporal objectives and identify which parts of a circuit are responsible for a given behavior. As with CoRa, however, corroborating CoRaDyn’s findings experimentally remains challenging, since constructing the locally analogous, feedback-disrupted system is not trivial. It may nonetheless be feasible, as suggested by Gómez-Schiavon & El-Samad [5]. Even so, this experimental caveat does not diminish CoRaDyn’s conceptual contribution.

The biological consequences of feedback control cannot be fully understood by asking only whether a system returns to its pre-perturbation state. It also requires asking how it gets there, and CoRaDyn provides a systematic way to ask that question.

## Data Availability

No experimental or externally generated datasets were used in this study. All results can be reproduced from the equations, parameter values, and algorithms provided in the Supplementary Information, together with the simulation and analysis code available in the GitHub repository https://github.com/SysEvo/CoRa_Dynamics.

## Code Availability

All code used to perform the simulations and analyses and to generate the figures presented in this study is freely available in the GitHub repository https://github.com/SysEvo/CoRa_Dynamics.

## Author Contributions

**Conceptualization:** M.G.-S.; **Methodology:** A.T.-L., M.G.-S.; **Software:** A.T.-L.; **Formal analysis:** A.T.-L.; **Investigation:** A.T.-L.; **Visualization:** A.T.-L., M.G.-S.; **Writing – original draft:** A.T.-L., M.G.-S.; **Writing – review & editing:** A.T.-L., M.G.-S.; **Supervision:** M.G.-S.; **Project administration:** M.G.-S.; **Funding acquisition:** M.G.-S.

## AI-use disclosure

Generative AI tools were used during manuscript preparation to assist with language editing, discussion of manuscript organization and presentation, and refinement of scientific communication. All scientific content, analyses, interpretations, and conclusions were developed and conducted by the authors. The manuscript was critically evaluated and approved by all authors, who take full responsibility for its accuracy and integrity.

## Acknowledgments

We thank Luis A. Aguilar from the *Laboratorio Nacional de Visualización Científica Avanzada* and Jair Santiago García Sotelo, Iliana Martínez, Rebeca Mucino, María Arteaga, and Eglee Lomelín from the *Laboratorio Internacional de Investigación sobre el Genoma Humano, Universidad Nacional Autónoma de México, Santiago de Querétaro, México*, for their support.

## Supporting information

**Supplementary Figures:** Fig S1-S16

**Supplementary Tables:** Table S1-S3

**Supplementary Text:** Section S1-S2

**Supplementary Code:** https://github.com/SysEvo/CoRa_Dynamics/releases/tag/v1.0.0.

## SUPPLEMENTARY INFORMATION

### S1 CoRaDyn approach

CoRaDyn builds on CoRa (Control Ratio), a method that quantifies how much feedback control contributes to a system’s ability to reject step perturbations (Gómez-Schiavon & El-Samad, 2022). Whereas CoRa evaluates feedback performance exclusively at steady state, CoRaDyn extends the framework to quantify feedback performance throughout the transient response, before steady state is reached. This provides complementary information by capturing dynamic features such as the magnitude of the initial deviation and the time required for the system to relax toward steady state.

To perform the analysis, both CoRa and CoRaDyn rely on a mathematically controlled comparison between two systems. One is the *feedback system*, which implements feedback control. The other is a *locally analogous system*, identical in structure and parameter values except for the removal of a single interaction that breaks the feedback loop under evaluation. This interaction is replaced with a constant input calibrated to reproduce the effect of the feedback at the steady state of the *feedback system* for the parameter set Θ.

The *locally analogous system* is constructed to ensure *internal equivalence*, such that all variables, biochemical reactions, parameter values, and interactions remain identical to those of the *feedback system*, except for the single interaction removed to disrupt the feedback mechanism under evaluation. It also ensures *external equivalence*, allowing both systems to converge to the same steady state. Consequently, any differences in the dynamics of the two systems following a small perturbation 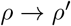, where *ρ* ∈ Θ, can be attributed exclusively to the functional contribution of the feedback mechanism.

To quantify these differences, CoRa compares the output of the *feedback* and *locally analogous systems* at a specified evaluation time, typically after both systems have reached steady state following a perturbation to *ρ*. Let *Y*_*ss*_ denote the steady-state value of the *feedback system*, and *Y*_*ss,F D*_ that of the *locally analogous system* for a given parameter set Θ. By construction, these values are identical, i.e., *Y*_*ss*_|_Θ_= *Y*_*ss,FD*_|_Θ_. Following a perturbation 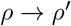, the corresponding steady-state outputs become 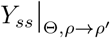 and 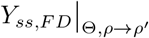, respectively. CoRa is then defined as:

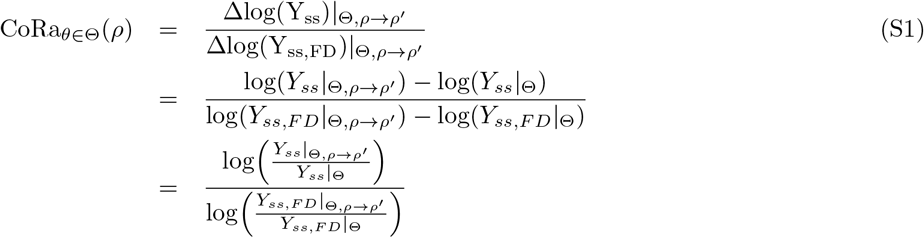

CoRaDyn, in contrast, quantifies the behavioral disparity between the *feedback* and *locally analogous systems* by integrating the deviation in the system output relative to its pre-perturbation state over time. Specifically, it evaluates this quantity from the time the perturbation is applied (*t*_0_) to a chosen evaluation time (*t*_*n*_), and compares the resulting cumulative deviation with that of the *locally analogous system*. To ensure a fair comparison, both integrals are normalized by the duration of the evaluation window (*t*_*n*_ − *t*_0_), although this normalization cancels in the final ratio. CoRaDyn is therefore defined as:

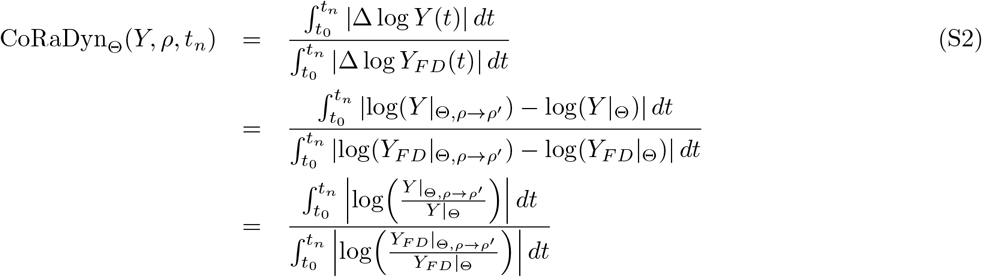

Here, *Y* |_Θ_ and *Y*_*F D*_|_Θ_ denote the output trajectories of the *feedback* and *locally analogous systems*, respectively, prior to the perturbation. Likewise, 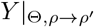 and 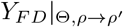 denote the corresponding trajectories following the perturbation 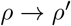, where *ρ* ∈ Θ.

Like CoRa, CoRaDyn yields a real number between 0 and 1. CoRaDyn_Θ_(*Y, ρ, t*_*n*_) approaches 0 when the feedback effectively attenuates the impact of the perturbation over the evaluation window, relative to the *locally analogous system*. This occurs when 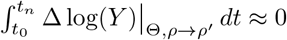, causing the ratio to become small. However, because CoRaDyn accumulates deviations throughout the evaluation window, transient dynamics occurring before the system reaches steady state contribute to the metric, even when perfect adaptation is ultimately achieved 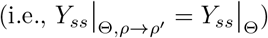. Consequently, systems exhibiting perfect adaptation generally yield positive CoRaDyn values over finite evaluation windows.

As CoRaDyn_Θ_(*Y, ρ, t*_*n*_) increases, it reflects a diminished contribution of feedback to perturbation rejection over the evaluation window. Nonetheless, as long as its value remains strictly between 0 and 1, feedback still reduces the effect of the perturbation relative to the *locally analogous system*, as reflected by the inequality

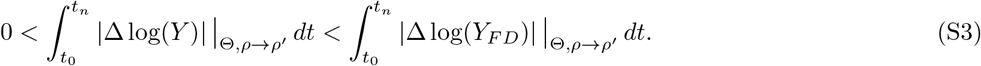

When CoRaDyn_Θ_(*Y, ρ, t*_*n*_) = 1, feedback has no effect on the system’s response over the evaluation window, making the output trajectory indistinguishable from that of the feedback-disrupted system:

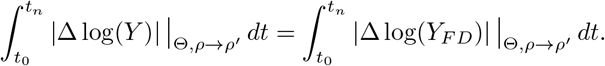

A defining feature of CoRaDyn is its dependence on *t*_*n*_, which determines the duration of the evaluation window and thereby influences the resulting value. When the system reaches a steady state after the perturbation 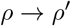, increasing *t*_*n*_ progressively reduces the relative contribution of transient dynamics. Consequently, the steady-state behavior dominates, and CoRaDyn converges to the corresponding CoRa value for the same parameter set Θ. Conversely, as *t*_*n*_ approaches *t*_0_, CoRaDyn approaches 1 because the outputs of the feedback and feedback-disrupted systems are initially nearly identical by construction. Only after sufficient time has elapsed (*t*_*n*_ *> t*_0_) does the corrective action of the feedback produce measurable differences between the trajectories of the two systems.

The choice of the evaluation time *t*_*n*_ is inherently context dependent and should be guided by the temporal characteristics of the biological system or mathematical model under study. For example, *t*_*n*_ may correspond to the time required for the system to reach a stable steady state or to settle into a limit cycle following a perturbation. Alternatively, it may be informed by empirical observations of the timescale over which a biological system typically mounts an effective corrective response. More generally, *t*_*n*_ should be chosen to reflect the timescales of the processes being evaluated. For instance, if the relevant dynamics occur over seconds, as in fast molecular interactions, selecting *t*_*n*_ on the order of seconds rather than minutes or hours ensures that the resulting CoRaDyn values capture the biologically relevant behavior.

In addition to the evaluation time, the perturbation magnitude is another important consideration when computing CoRaDyn. Whereas CoRa is approximately independent of perturbation size, provided the perturbation remains sufficiently small, CoRaDyn can be sensitive to it. This is because CoRaDyn captures features of the system’s full dynamical response, including response amplitude, adaptation time, and transient excursions, all of which may vary with perturbation magnitude. For example, in systems exhibiting damped oscillations, larger perturbations can prolong transients or amplify the output response, thereby altering the CoRaDyn value (Fig. S1). Furthermore, this sensitivity depends not only on the system’s dynamical regime but also on the evaluation time *t*_*n*_. Shorter evaluation windows place greater emphasis on early-time dynamics, which are generally more sensitive to perturbation magnitude. In addition, sufficiently large perturbations may drive the system outside the regime in which the feedback and feedback-disrupted systems remain locally analogous, introducing differences that are no longer attributable solely to the presence or absence of feedback. Therefore, small perturbations are generally preferred to preserve local analogy and ensure a meaningful comparison. Nevertheless, the appropriate perturbation magnitude should ultimately be determined on a case-by-case basis by assessing its influence on both the system dynamics and the resulting CoRaDyn value.

Finally, interpreting CoRaDyn values requires careful consideration. Lower values do not necessarily indicate better system performance, since different dynamic responses may be advantageous depending on the physiological context and functional objectives of the system. For example, when the goal is to maintain an output close to its pre-perturbation level throughout the evaluation window, lower CoRaDyn values are generally desirable. In contrast, systems that rely on transient responses, such as pulsatile signaling or delayed adaptation, may naturally exhibit higher CoRaDyn values while still achieving their intended function. The key question is whether feedback improves system behavior relative to its feedback-disrupted counterpart with respect to the biological function under consideration. Values approaching one indicate that feedback contributes little to the system’s response, as the feedback and feedback-disrupted systems behave similarly over the evaluation window. Beyond this, no universal threshold exists; the interpretation of CoRaDyn values ultimately depends on the physiological constraints and functional objectives of the system.

### S2 Biomolecular PID controller as a case study

#### Adaptation of the PID system proposed by Chevalier et al. (2019)

The model proposed by Chevalier et al. (2019) extends the antithetic integral feedback motif introduced by Briat et al. [18] to implement modular proportional, integral, and derivative (PID) control.

The core network comprises four molecular species connected through three regulatory interactions. At its center is the antithetic motif, formed by the controller species *Z*_1_ and *Z*_2_, which undergo a mutual annihilation reaction. This interaction sequesters both controller species through stoichiometric binding, thereby implementing the integral feedback mechanism. In this architecture, *Z*_1_ activates the production of *X*_1_, which in turn drives the production of the system output, *X*_*C*_. The output *X*_*C*_ subsequently induces the production of *Z*_2_, thereby closing the feedback loop. As a result, the feedback maintains *X*_*C*_ at its target level despite persistent disturbances.

The dynamics of this PID feedback system are described by the following set of differential equations:

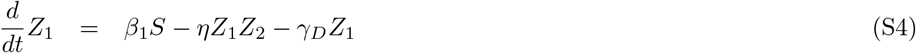

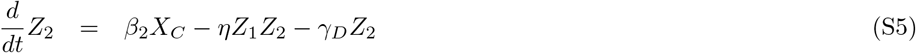

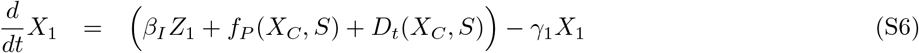

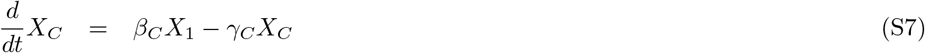

Here, *S* represents the input signal; *β*_*I*_ *Z*_1_ introduces **integral control** through the antithetic feedback motif; *f*_*P*_ (*X*_*C*_, *S*) represents **proportional control** (Eq. S8); and *D*_*t*_(*X*_*C*_, *S*) represents **derivative control** (Eqs. S10–S11). Together, these terms integrate integral, proportional, and derivative feedback into a unified PID control architecture that combines robust perfect adaptation with tunable transient dynamics.

**Proportional control** is implemented through an inhibitory function *f*_*P*_ (*X*_*C*_, *S*) that modulates the production of *X*_1_ in response to deviations in *X*_*C*_. Specifically, this term is defined as:

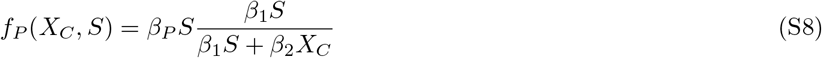

This formulation enables the strength of *X*_1_ production to vary proportionally with the difference between the desired and actual output levels, thereby contributing to proportional feedback control.

The **derivative control** component is implemented through a two-node dynamical module that approximates the time derivative of *X*_*C*_. This module consists of two additional species, *A* and *M*, whose dynamics are described by the following equations:

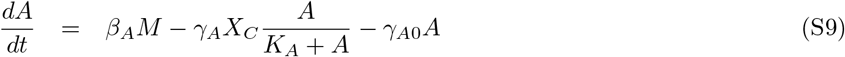

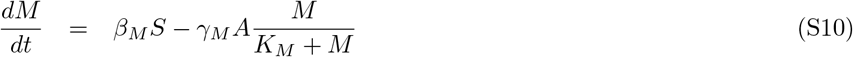

The derivative term is then defined as

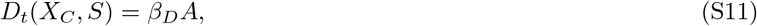

where *A* provides a dynamical approximation of the temporal changes in *X*_*C*_, enabling the system to adjust its response based on derivative feedback.

#### Locally analogous feedback-disrupted systems for the PID system

To evaluate how PID control contributes to the system’s transient dynamics and disturbance rejection capacity using CoRaDyn, we construct the corresponding locally analogous feedback-disrupted systems. Because the PID controller comprises three distinct feedback components—integral, proportional, and derivative control—we construct component-specific feedback-disrupted systems in which only the interaction associated with the feedback mechanism under evaluation is disrupted. In addition, we construct a fully feedback-disrupted system in which all three feedback loops are simultaneously disrupted, providing a reference for evaluating the overall contribution of PID control.

To disrupt the contribution of all three feedback components, we remove the interaction between *X*_*C*_ and *Z*_2_ that closes the antithetic integral feedback loop. The fully feedback-disrupted system is obtained by replacing Eq. S6 with:

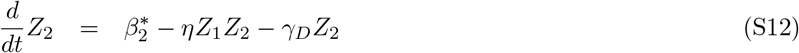

where *β*^*^ = *β*_2_*X*_*C,ss*_ is calibrated to preserve the pre-perturbation steady state.

To disrupt individual feedback components, we instead modify Eq. S7. When proportional feedback is disrupted, the dynamic proportional term *f*_*P*_ (*X*_*C*_, *S*) is replaced by its steady-state equivalent:

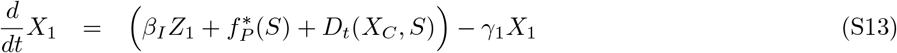

where

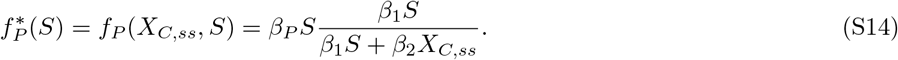

Similarly, when derivative feedback is disrupted, the dynamic derivative term *D*_*t*_(*X*_*C*_, *S*) is replaced by its steady-state equivalent:

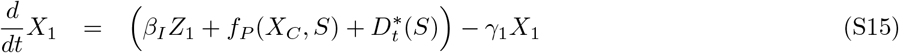

where

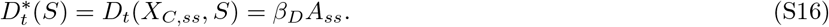

Finally, to disrupt both proportional and derivative feedback, both dynamic terms are replaced by their steady-state equivalents:

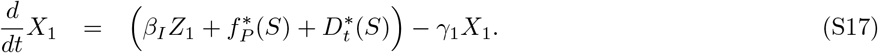

**Fig S1.**
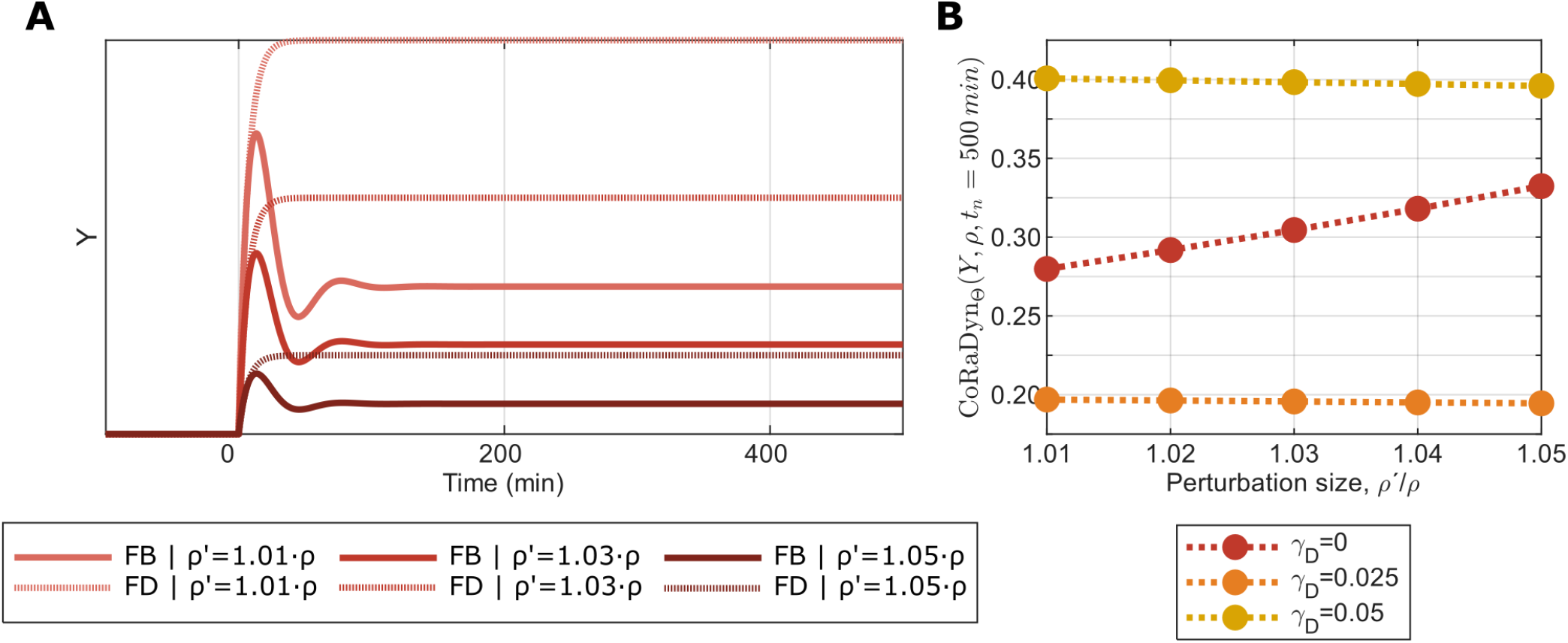
Effect of perturbation magnitude on CoRaDyn values. **A:** CoRaDyn as a function of the evaluation time (*t*_*n*_) for an example system subjected to three different perturbation magnitudes, 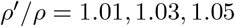. **B:** CoRaDyn evaluated at *t*_*n*_ = 500 min for a broader set of perturbation magnitudes spanning the same range illustrated in **A** and for three parameterizations of the system (*γ*_*D*_ = 0, red, as shown in **A**; *γ*_*D*_ = 0.025, orange; and *γ*_*D*_ = 0.05, yellow). While CoRa is approximately independent of perturbation magnitude for sufficiently small perturbations, CoRaDyn may depend on perturbation magnitude, with the degree of sensitivity determined by the system dynamics and parameterization.

**Table S1.** Capabilities of different frameworks for evaluating biological feedback control. Comparison of adaptation time, CoRa, and CoRaDyn with respect to the types of feedback behavior they can evaluate and the questions they address.

| Capability | Adaptation Time<sup>a</sup> | CoRa<sup>b</sup> | CoRaDyn |
| --- | --- | --- | --- |
| Quantifies steady-state adaptation | — | ✓ | ✓<sup>c</sup> |
| Quantifies transient feedback performance | Limited | — | ✓ |
| Defined for sustained post-perturbation oscillations | — | — | ✓ |
| Distinguishes controllers with identical steady-state adaptation | — | — | ✓ |
| Allows evaluation at arbitrary temporal scales | — | — | ✓ |
| Isolates the contribution of individual feedback loops through controlled comparison | — | Proposed<sup>d</sup> | ✓ |
| Requires a stable pre-perturbation steady state | ✓ | ✓ | ✓ |

<sup>a</sup> As proposed in Chevalier et al. (2019).
<sup>b</sup> As proposed in Gómez-Schiavon & El-Samad (2022).
<sup>c</sup> CoRaDyn converges to CoRa in the long-time limit for systems that reach a stable post-perturbation steady state.
<sup>d</sup> The use of locally analogous systems to isolate individual feedback loops was discussed in the original CoRa formulation, but was not explicitly demonstrated.

**Fig S2.**
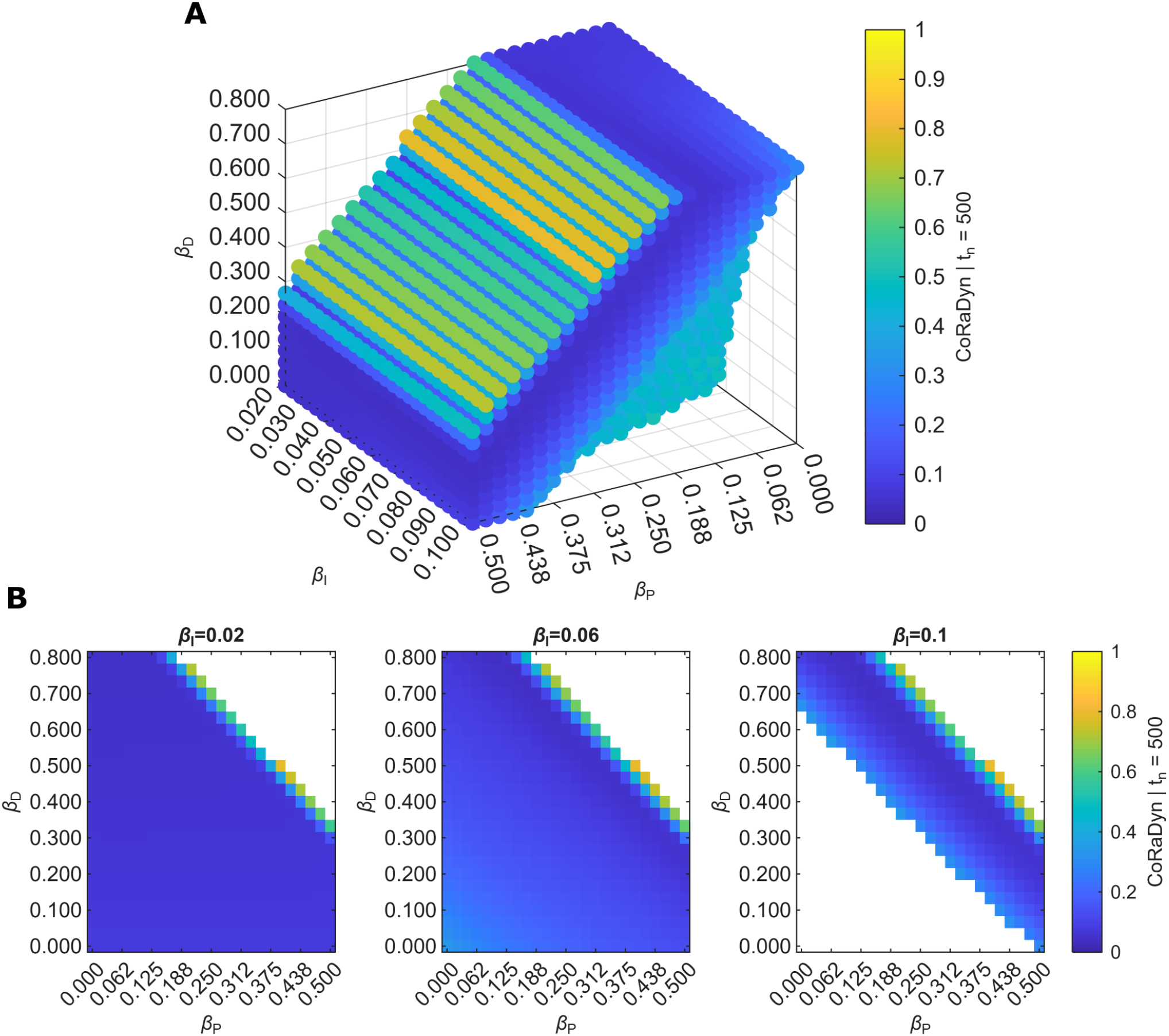
CoRaDyn evaluated at a fixed time reproduces the parameter dependence of PID control performance. CoRaDyn evaluated at a representative evaluation time (*t*_*n*_ = 500 min) over the proportional (*β*_*P*_), integral (*β*_*I*_), and derivative (*β*_*D*_) controller weights. **A:** CoRaDyn across the complete explored parameter space. Axes correspond to the proportional (*β*_*P*_), integral (*β*_*I*_), and derivative (*β*_*D*_) control weights. **B:** Two-dimensional slices of the parameter space for representative values of the integral control weight (*β*_*I*_ = 0.02, 0.06, and 0.10 min^*−*1^; indicated above each panel). Colors indicate the CoRaDyn value according to the color scale. White regions indicate parameter combinations for which no stable pre-perturbation steady state exists and therefore CoRaDyn cannot be evaluated. See *Section S2* for equations and Table S3 for parameter values.

**Fig S3.**
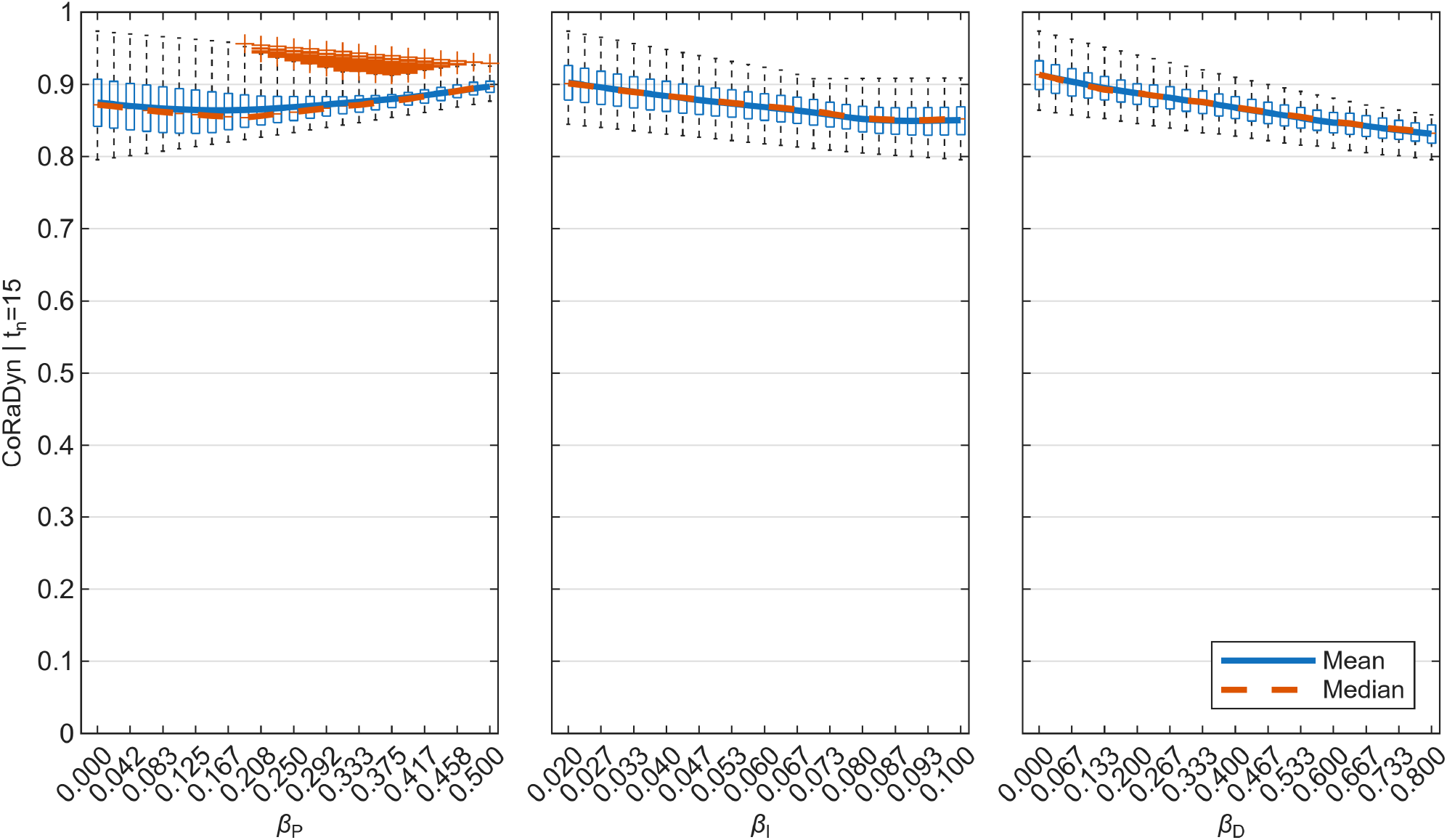
Distribution of CoRaDyn values across PID controller weights at *t*_*n*_ = 15 min. Boxplots of CoRaDyn for fixed values of the proportional (*β*_*P*_; left), integral (*β*_*I*_; center), and derivative (*β*_*D*_; right) control weights. For each control weight value, the distribution is computed over all combinations of the remaining two controller weights. Blue lines denote the mean and red lines the median. See *Section S2* for equations and Table S3 for parameter values.

**Fig S4.**
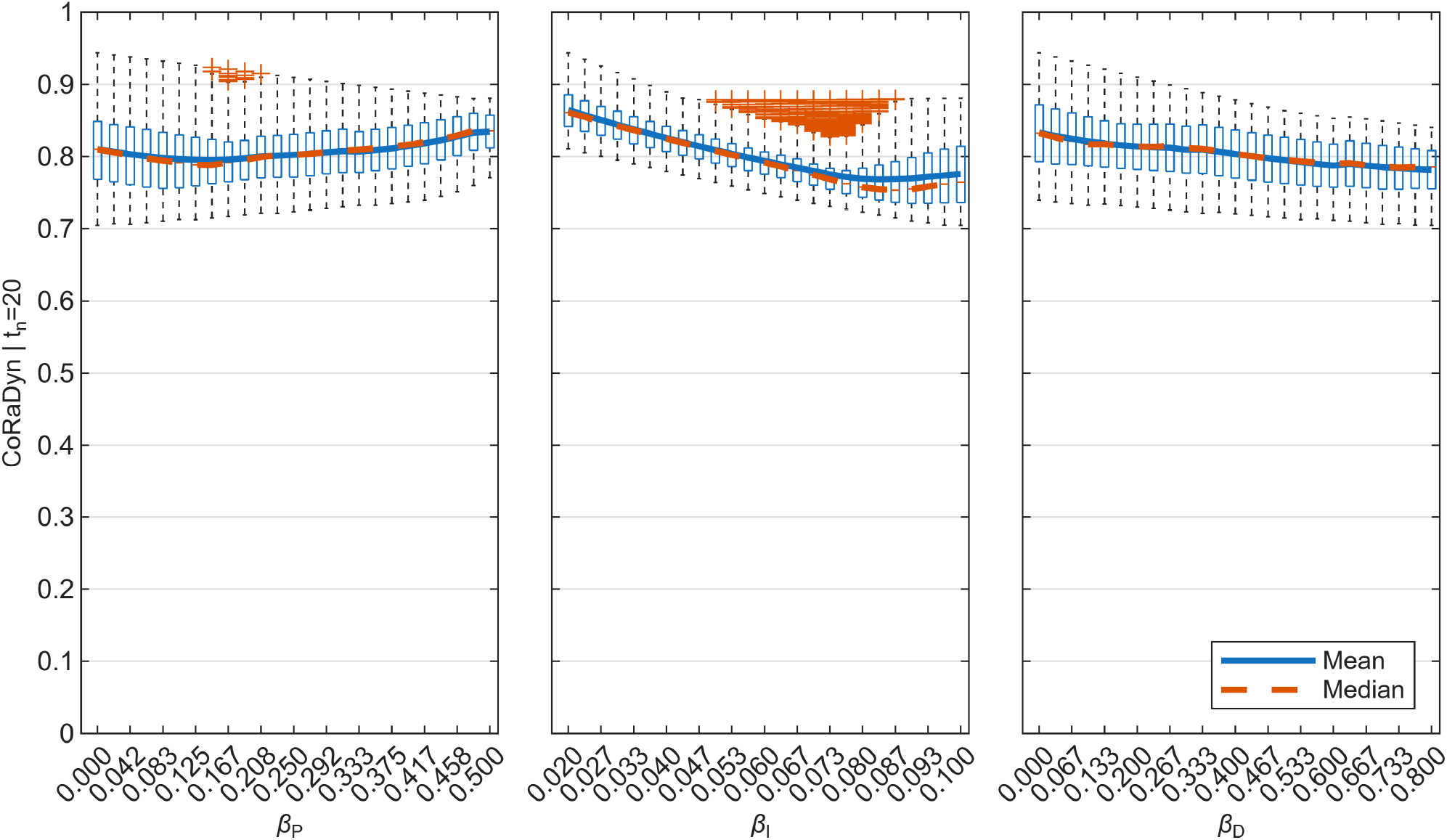
Distribution of CoRaDyn values across PID controller weights at *t*_*n*_ = 20 min. Boxplots of CoRaDyn for fixed values of the proportional (*β*_*P*_; left), integral (*β*_*I*_; center), and derivative (*β*_*D*_; right) control weights. For each control weight value, the distribution is computed over all combinations of the remaining two controller weights. Blue lines denote the mean and red lines the median. See *Section S2* for equations and Table S3 for parameter values.

**Fig S5.**
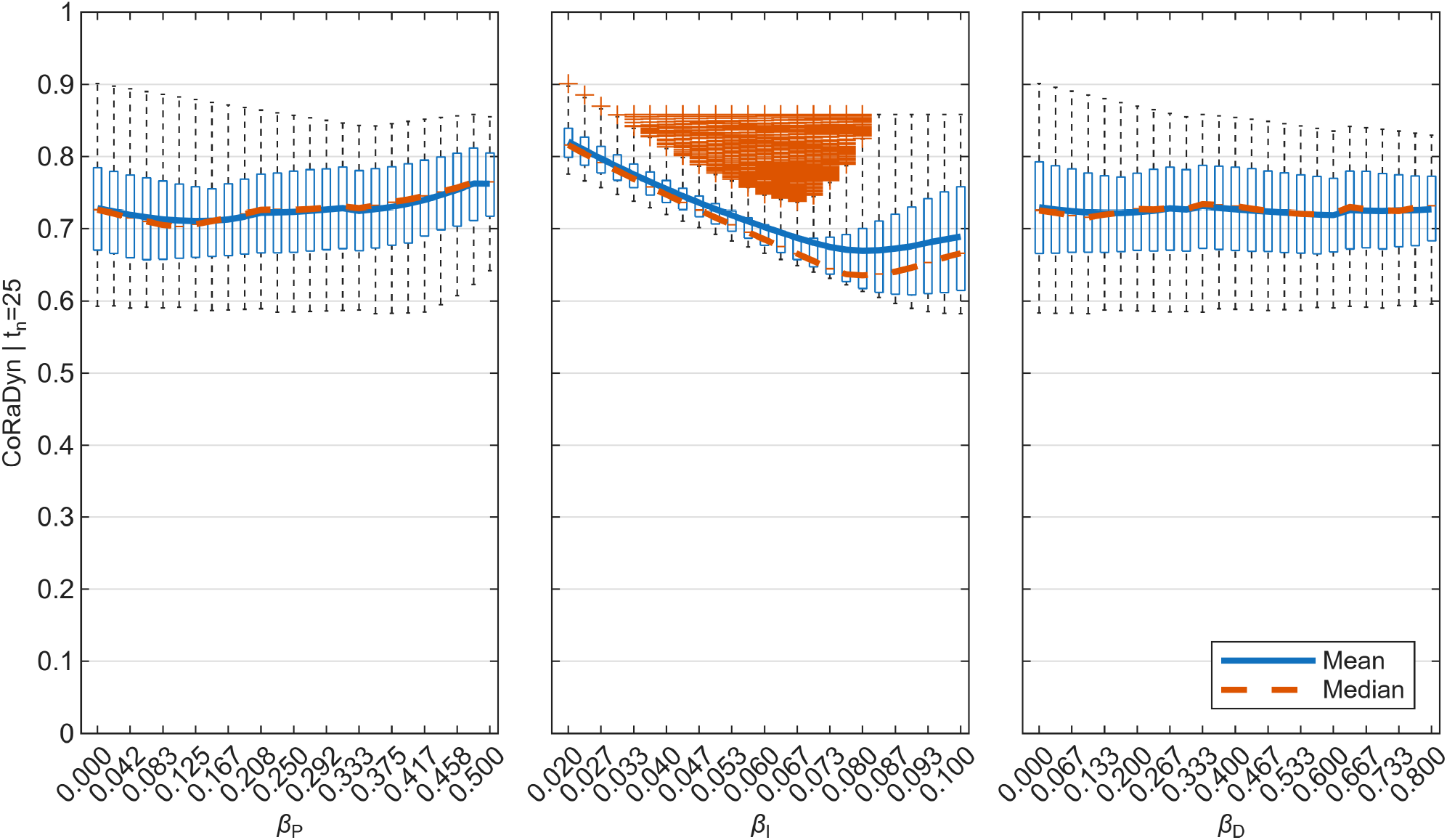
Distribution of CoRaDyn values across PID controller weights at *t*_*n*_ = 25 min. Boxplots of CoRaDyn for fixed values of the proportional (*β*_*P*_; left), integral (*β*_*I*_; center), and derivative (*β*_*D*_; right) control weights. For each control weight value, the distribution is computed over all combinations of the remaining two controller weights. Blue lines denote the mean and red lines the median. See *Section S2* for equations and Table S3 for parameter values.

**Fig S6.**
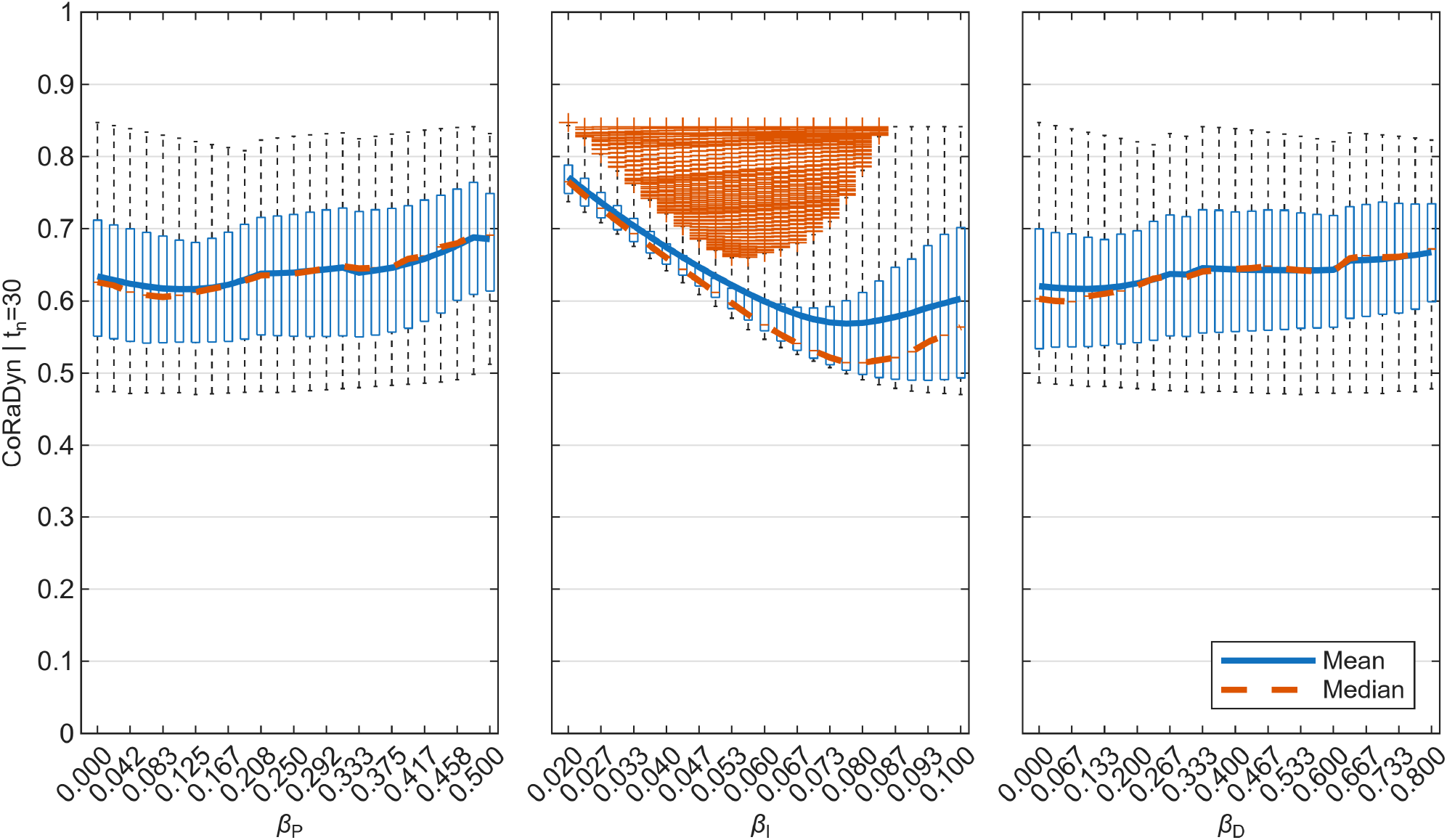
Distribution of CoRaDyn values across PID controller weights at *t*_*n*_ = 30 min. Boxplots of CoRaDyn for fixed values of the proportional (*β*_*P*_; left), integral (*β*_*I*_; center), and derivative (*β*_*D*_; right) control weights. For each control weight value, the distribution is computed over all combinations of the remaining two controller weights. Blue lines denote the mean and red lines the median. See *Section S2* for equations and Table S3 for parameter values.

**Fig S7.**
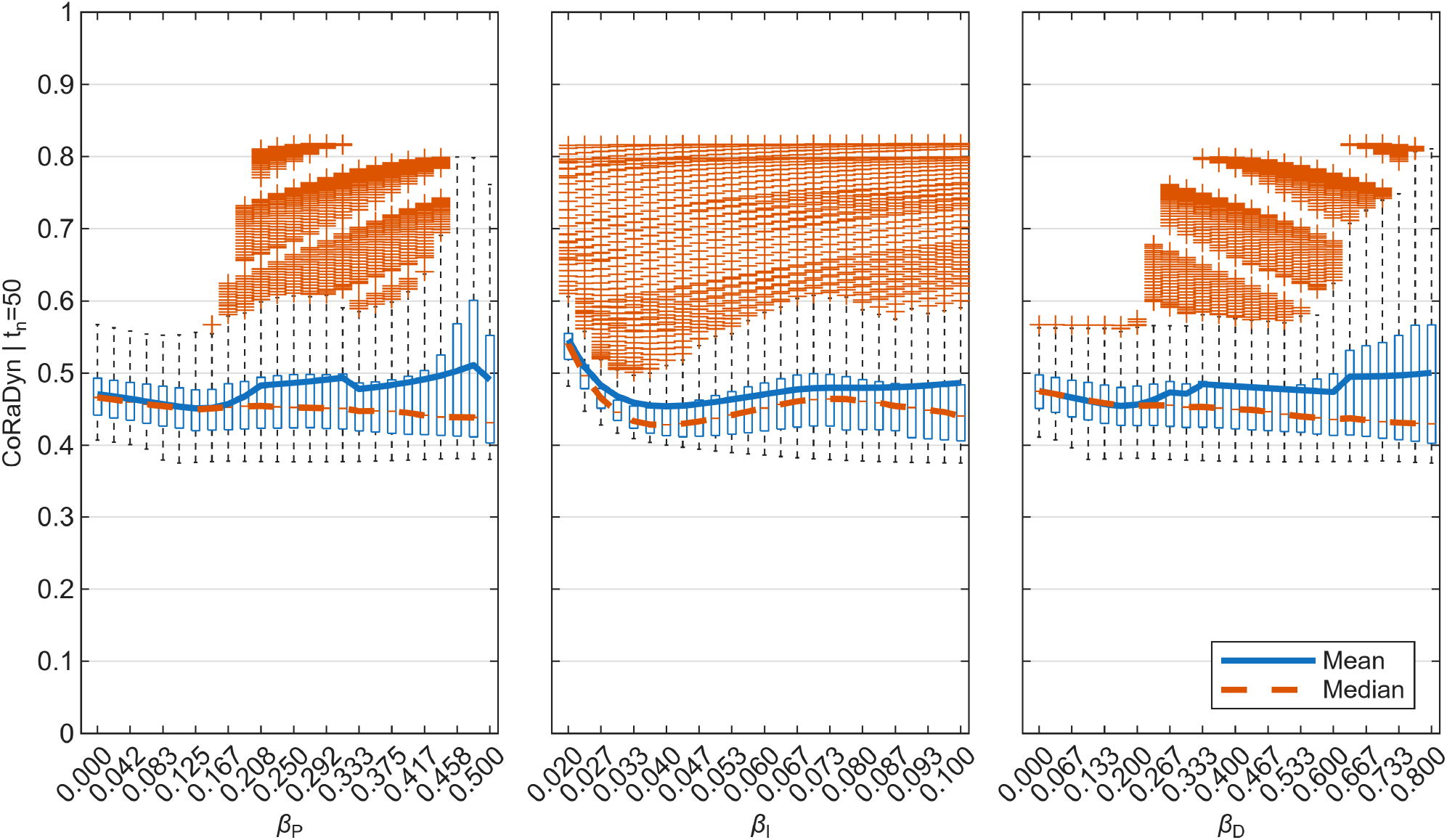
Distribution of CoRaDyn values across PID controller weights at *t*_*n*_ = 50 min. Boxplots of CoRaDyn for fixed values of the proportional (*β*_*P*_; left), integral (*β*_*I*_; center), and derivative (*β*_*D*_; right) control weights. For each control weight value, the distribution is computed over all combinations of the remaining two controller weights. Blue lines denote the mean and red lines the median. See *Section S2* for equations and Table S3 for parameter values.

**Fig S8.**
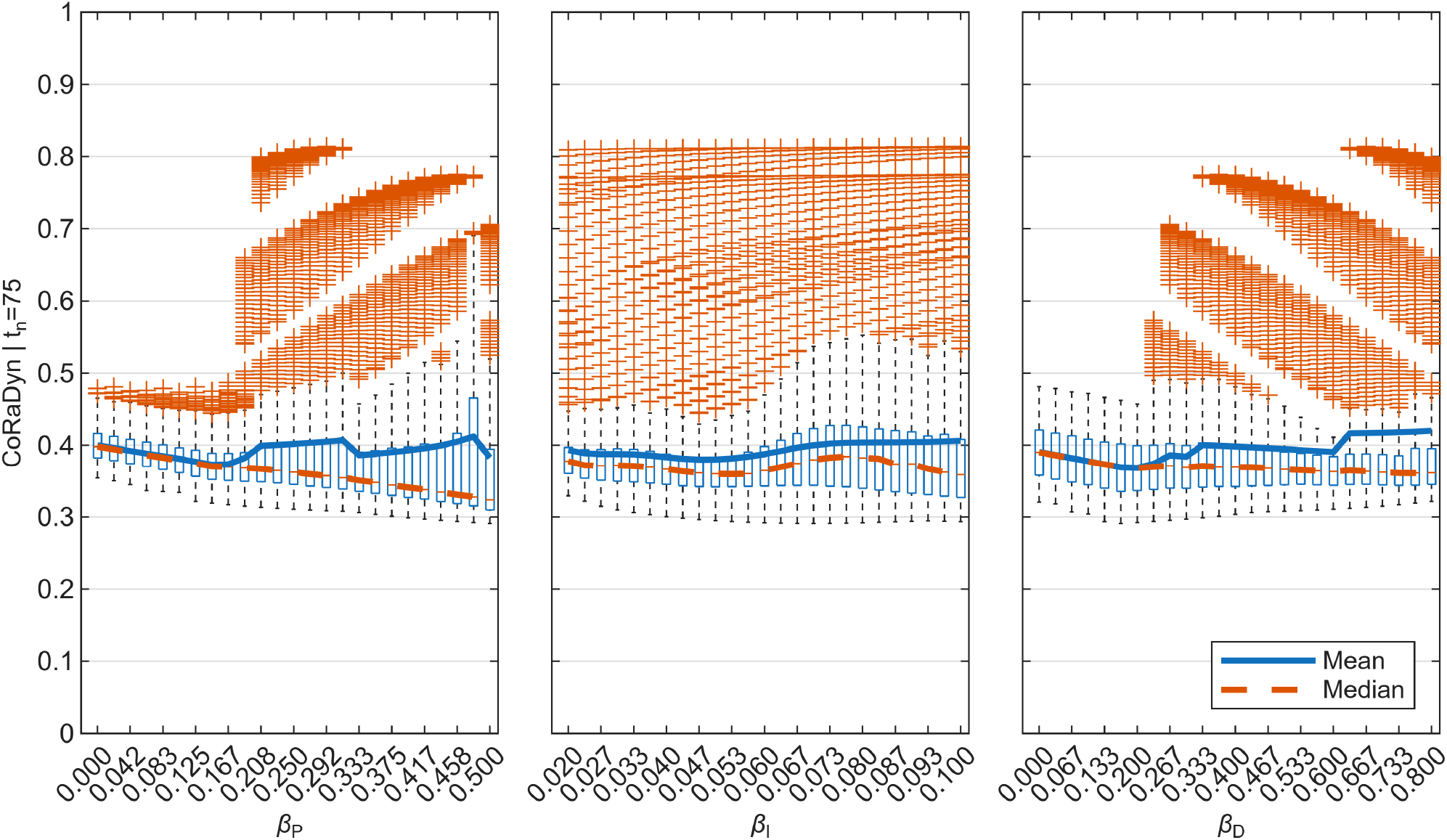
Distribution of CoRaDyn values across PID controller weights at *t*_*n*_ = 75 min. Boxplots of CoRaDyn for fixed values of the proportional (*β*_*P*_; left), integral (*β*_*I*_; center), and derivative (*β*_*D*_; right) control weights. For each control weight value, the distribution is computed over all combinations of the remaining two controller weights. Blue lines denote the mean and red lines the median. See *Section S2* for equations and Table S3 for parameter values.

**Fig S9.**
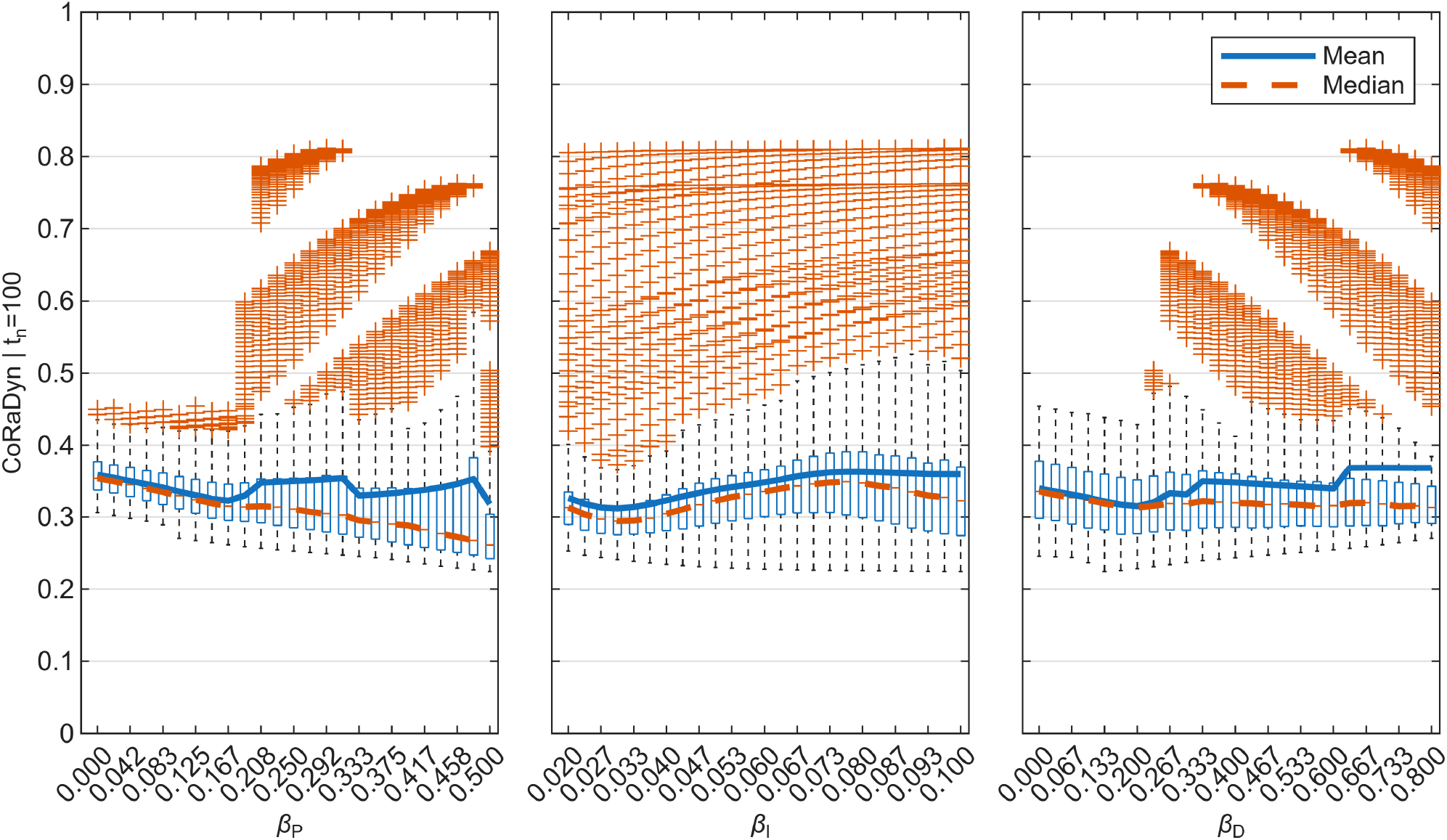
Distribution of CoRaDyn values across PID controller weights at *t*_*n*_ = 100 min. Boxplots of CoRaDyn for fixed values of the proportional (*β*_*P*_; left), integral (*β*_*I*_; center), and derivative (*β*_*D*_; right) control weights. For each control weight value, the distribution is computed over all combinations of the remaining two controller weights. Blue lines denote the mean and red lines the median. See *Section S2* for equations and Table S3 for parameter values.

**Fig S10.**
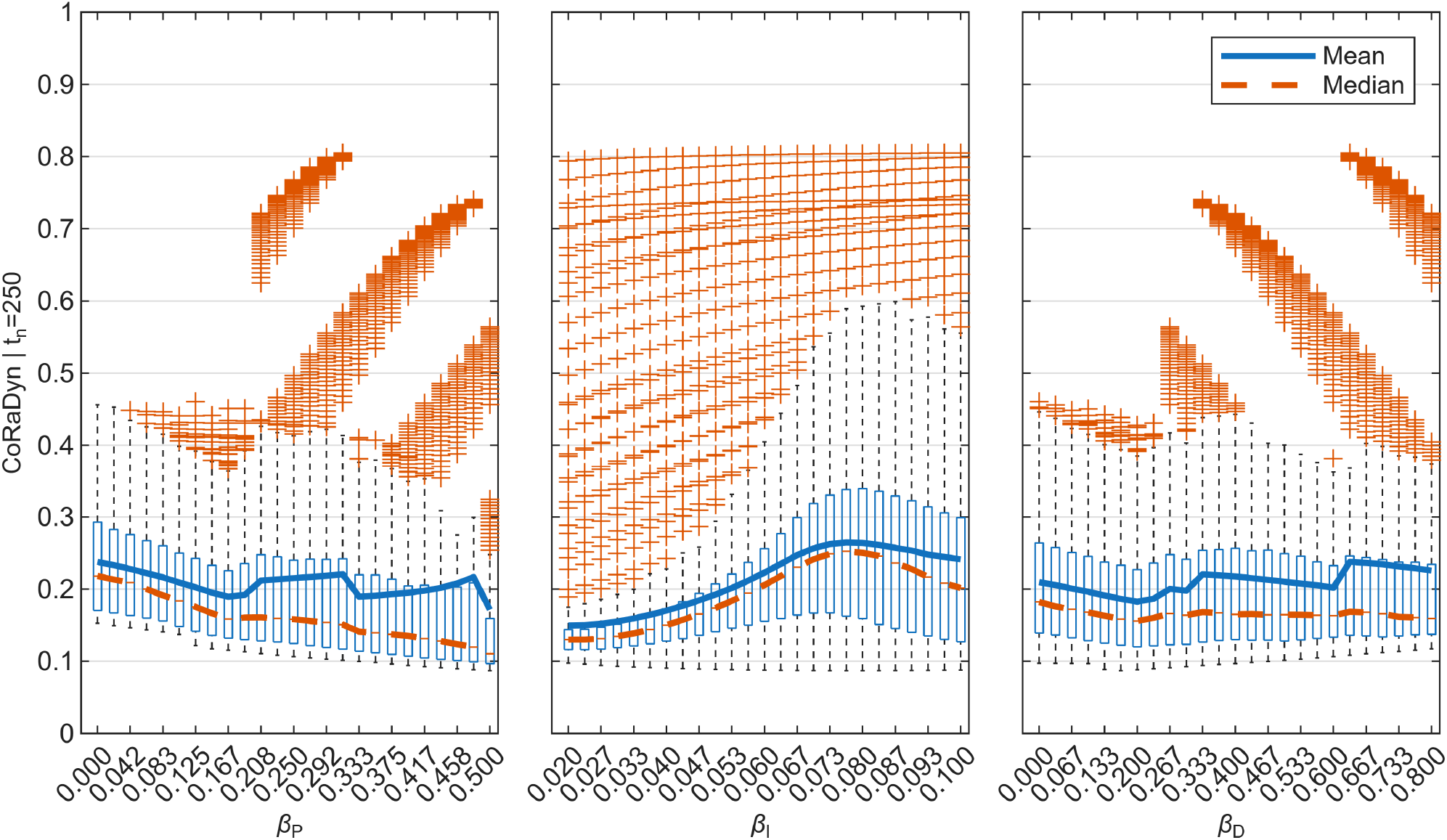
Distribution of CoRaDyn values across PID controller weights at *t*_*n*_ = 250 min. Boxplots of CoRaDyn for fixed values of the proportional (*β*_*P*_; left), integral (*β*_*I*_; center), and derivative (*β*_*D*_; right) control weights. For each control weight value, the distribution is computed over all combinations of the remaining two controller weights. Blue lines denote the mean and red lines the median. See *Section S2* for equations and Table S3 for parameter values.

**Fig S11.**
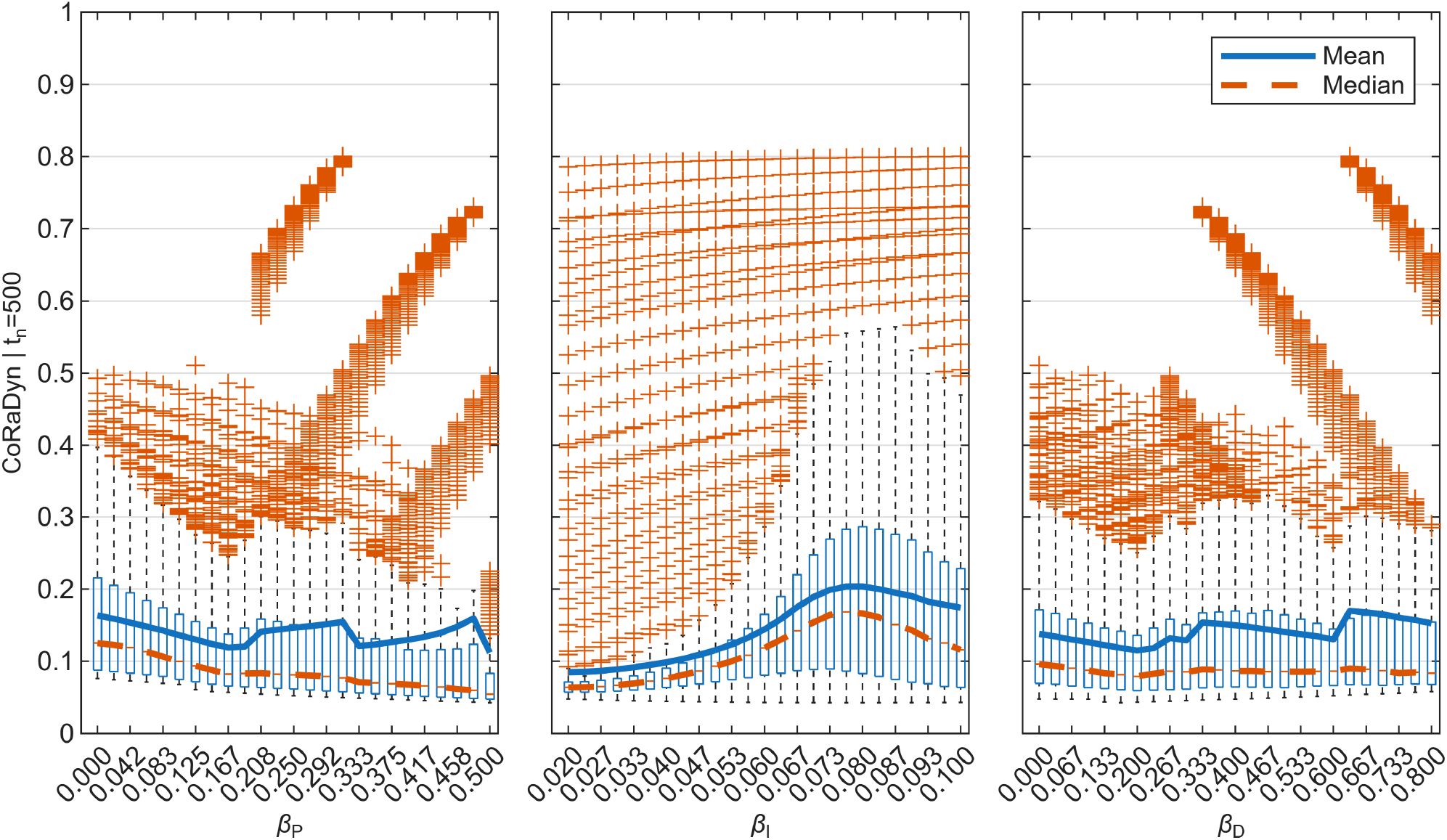
Distribution of CoRaDyn values across PID controller weights at *t*_*n*_ = 500 min. Boxplots of CoRaDyn for fixed values of the proportional (*β*_*P*_; left), integral (*β*_*I*_; center), and derivative (*β*_*D*_; right) control weights. For each control weight value, the distribution is computed over all combinations of the remaining two controller weights. Blue lines denote the mean and red lines the median. See *Section S2* for equations and Table S3 for parameter values.

**Fig S12.**
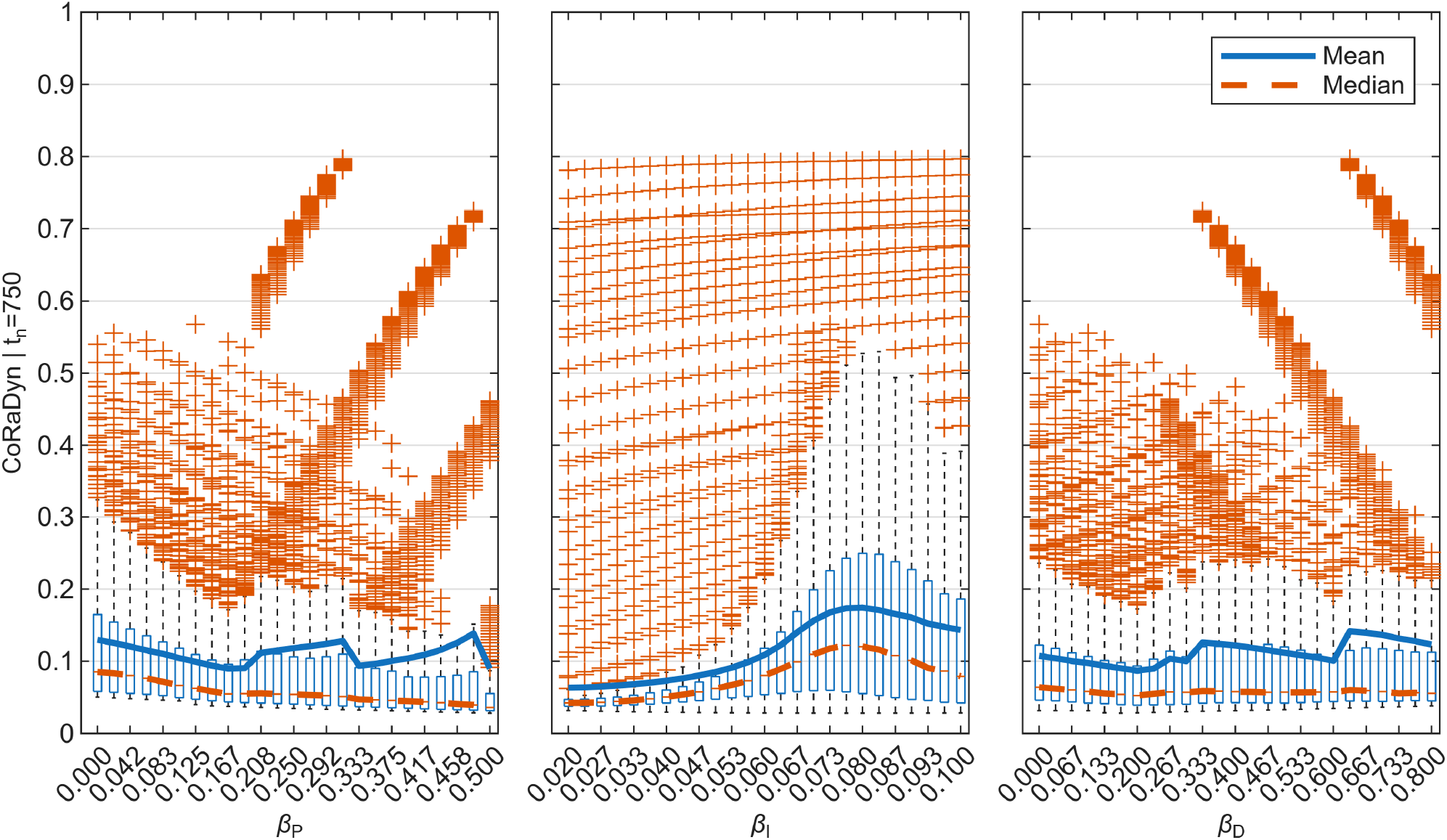
Distribution of CoRaDyn values across PID controller weights at *t*_*n*_ = 750 min. Boxplots of CoRaDyn for fixed values of the proportional (*β*_*P*_; left), integral (*β*_*I*_; center), and derivative (*β*_*D*_; right) control weights. For each control weight value, the distribution is computed over all combinations of the remaining two controller weights. Blue lines denote the mean and red lines the median. See *Section S2* for equations and Table S3 for parameter values.

**Fig S13.**
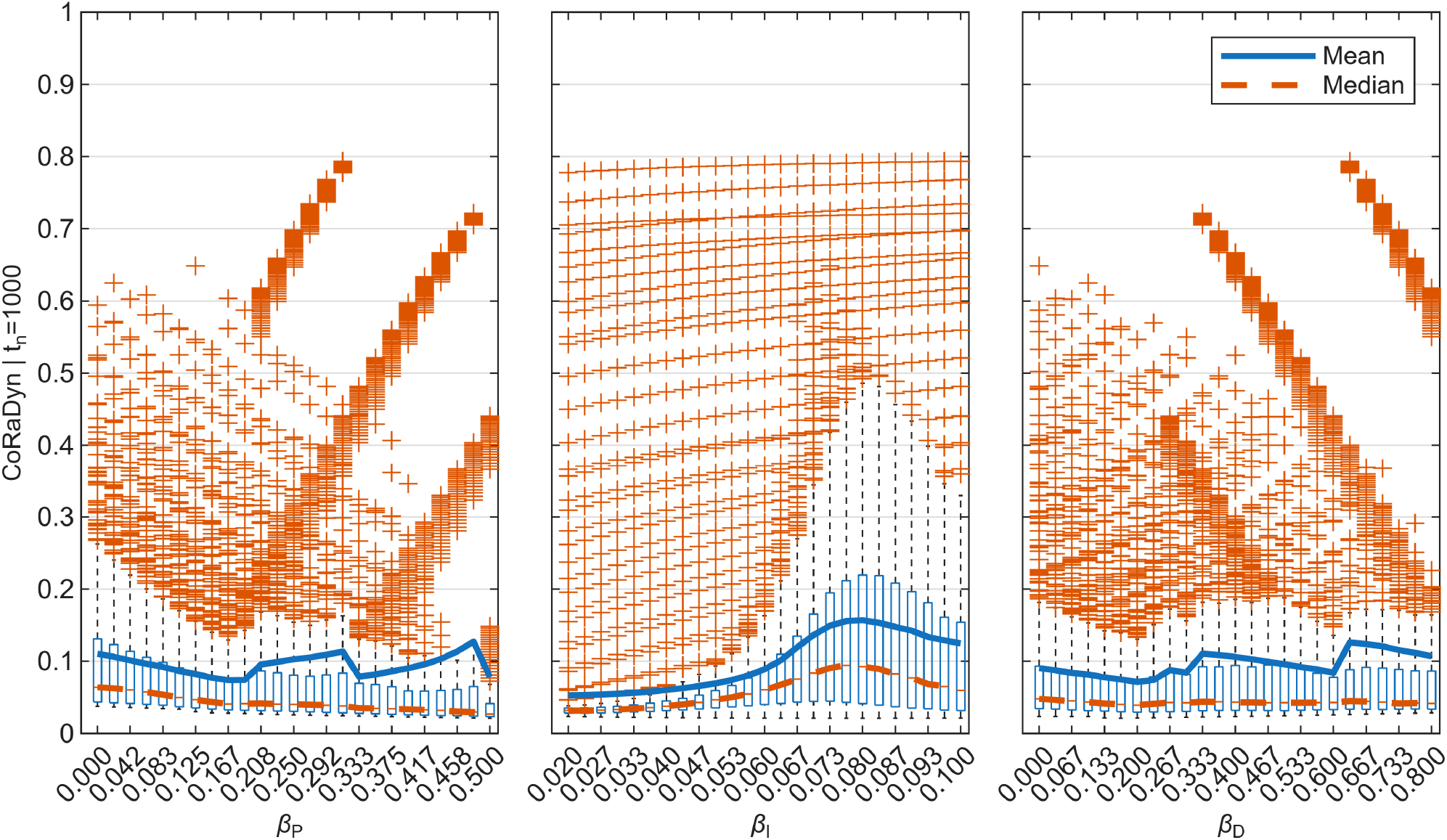
Distribution of CoRaDyn values across PID controller weights at *t*_*n*_ = 1000 min. Boxplots of CoRaDyn for fixed values of the proportional (*β*_*P*_; left), integral (*β*_*I*_; center), and derivative (*β*_*D*_; right) control weights. For each control weight value, the distribution is computed over all combinations of the remaining two controller weights. Blue lines denote the mean and red lines the median. See *Section S2* for equations and Table S3 for parameter values.

**Fig S14.**
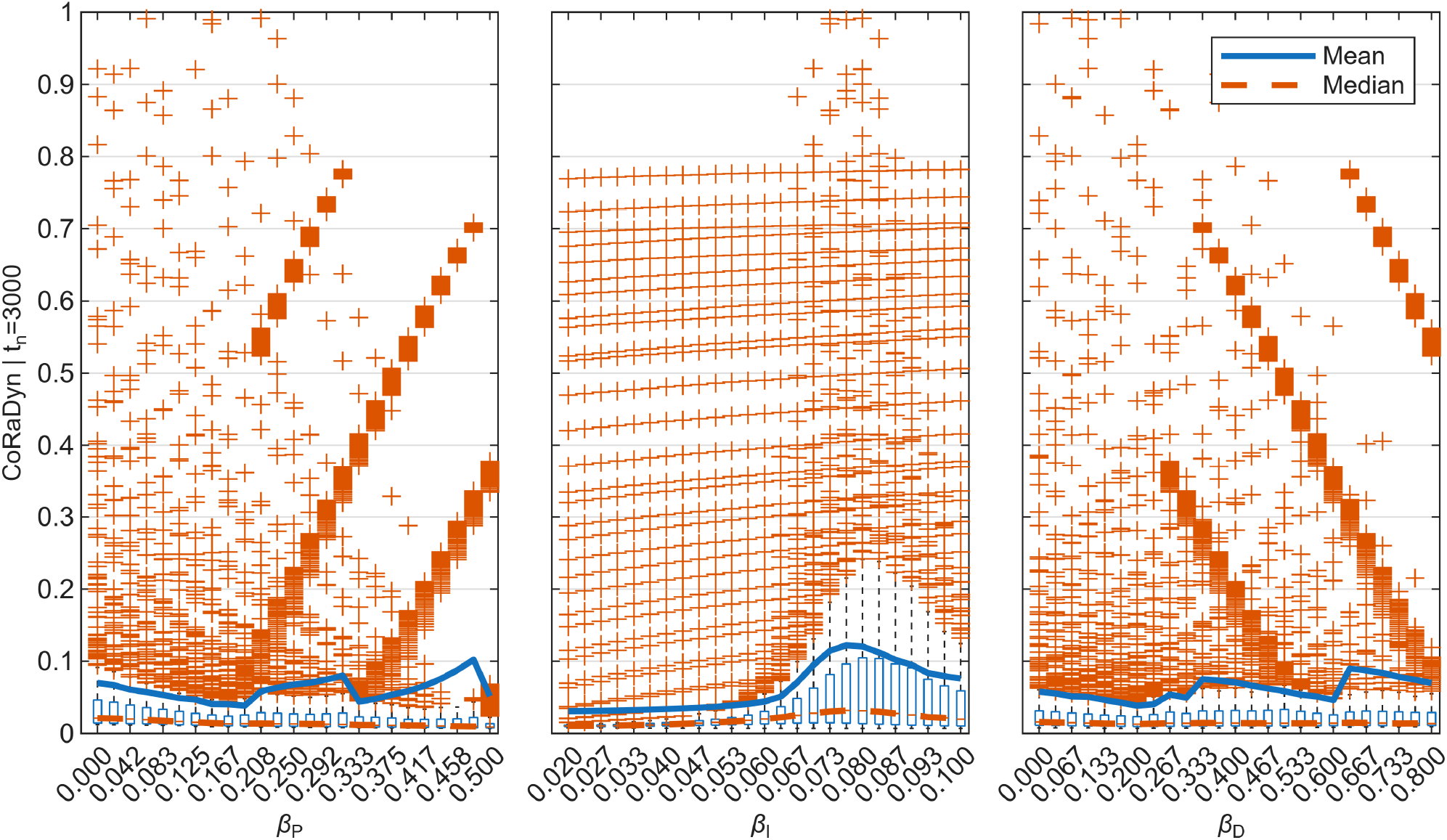
Distribution of CoRaDyn values across PID controller weights at *t*_*n*_ = 3000 min. Boxplots of CoRaDyn for fixed values of the proportional (*β*_*P*_; left), integral (*β*_*I*_; center), and derivative (*β*_*D*_; right) control weights. For each control weight value, the distribution is computed over all combinations of the remaining two controller weights. Blue lines denote the mean and red lines the median. See *Section S2* for equations and Table S3 for parameter values.

**Fig S15.**
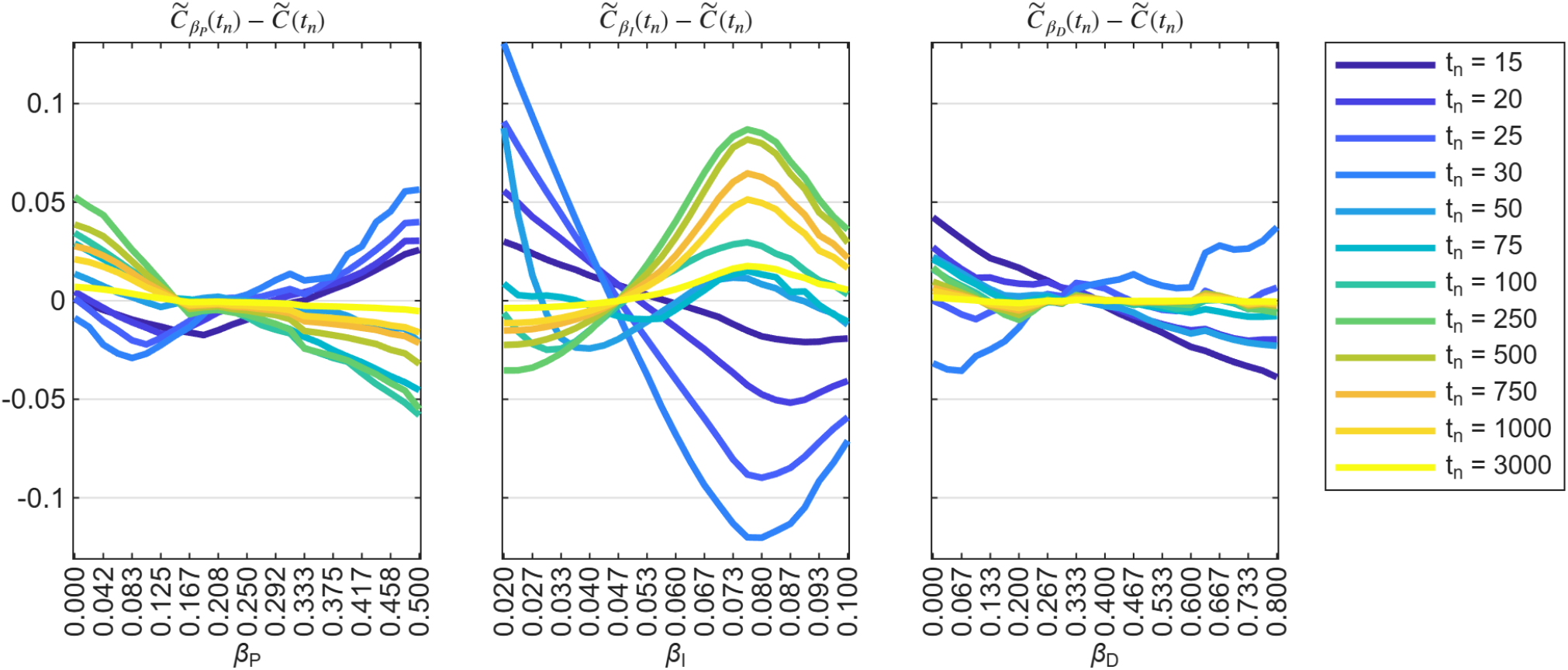
Deviations from the global median CoRaDyn across evaluation times. Differences between the median CoRaDyn associated with each controller weight and the corresponding global median, 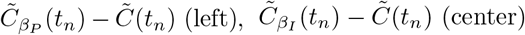, and 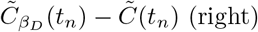, shown as functions of the corresponding controller weight. Within each panel, colored curves represent different evaluation times (*t*_*n*_; see legend). This representation facilitates comparison of how the influence of each controller weight evolves throughout the adaptive response. See *Section S2* for equations and Table S3 for parameter values.

**Fig S16.**
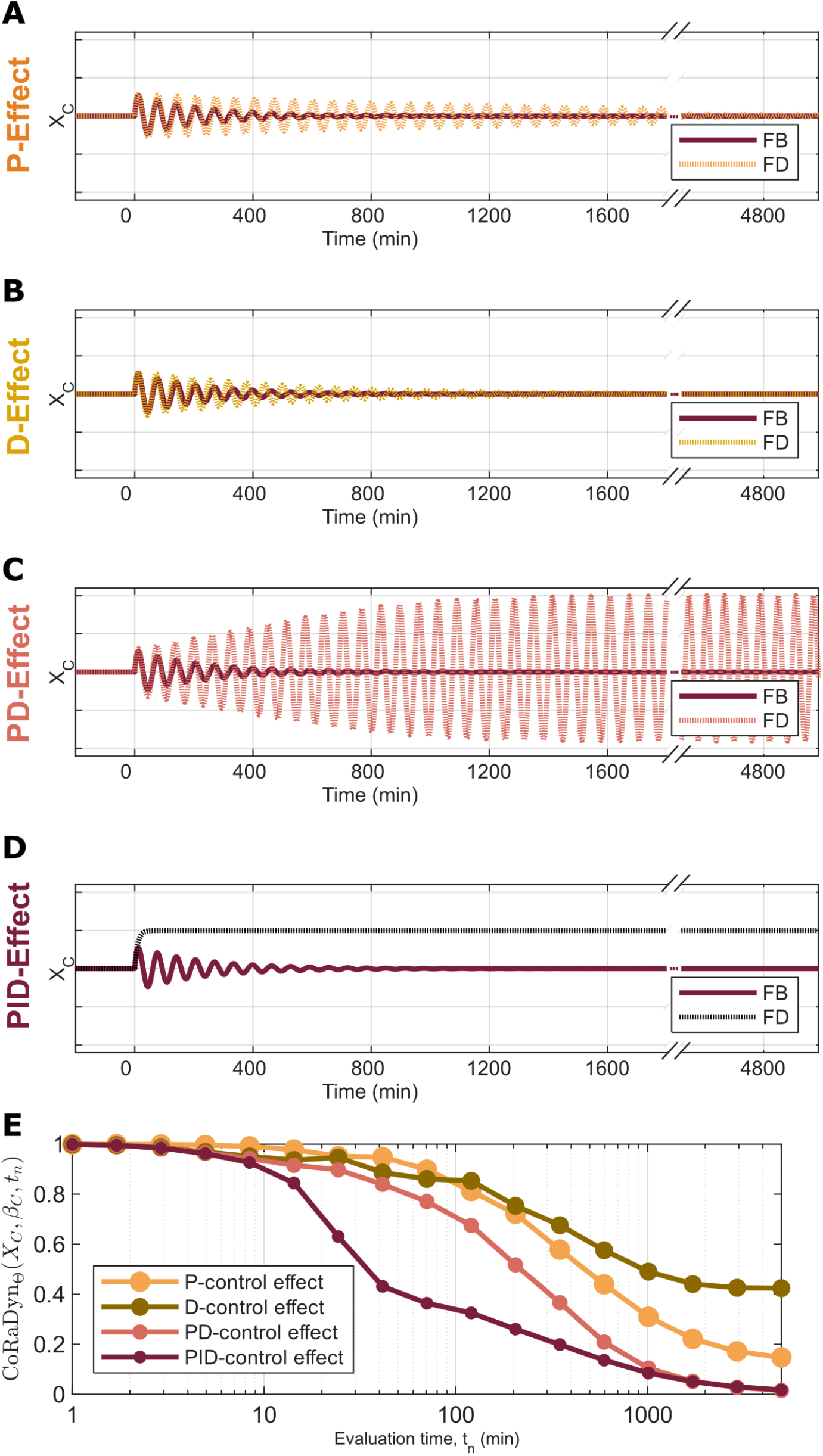
CoRaDyn isolates the dynamic contribution of overlapping feedback loops under an alternative parameterization. **A–D:** Dynamic response of the controlled variable (*X*_*C*_) following a step perturbation over *β*_*C*_ ∈ Θ applied at time zero. In each panel, the original PID controller (dark red) is compared with its corresponding locally analogous system, in which selected feedback components have been disrupted while preserving the remaining network unchanged: (**A**) proportional feedback removed (orange), (**B**) derivative feedback removed (yellow), (**C**) proportional and derivative feedback removed (red), and (**D**) all feedback components removed (black). **E:** CoRaDyn as a function of the evaluation time (*t*_*n*_) for the controlled comparisons shown in panels A–D, using the same color scheme. As in Fig. 5, the locally analogous systems in panels A–C retain integral control and therefore exhibit perfect adaptation despite their altered transient responses. Consequently, CoRaDyn quantifies the cumulative dynamic contribution of the disrupted feedback component(s), enabling the individual and combined effects of proportional, derivative, and integral control to be compared within the same controller architecture. This alternative parameterization illustrates that the quantitative contribution of each feedback component depends on the underlying system parameters. See *Section S2* for equations and Table S3 for parameter values.

**Table S2.** Used parameter values in main figures.

| Figure | CoRaDyn parameters | Model parameters |
| --- | --- | --- |
| **Fig. 2A** | Model: Eqs. S5 - S7<br>Controlled variable, $Y: X_C$<br>Perturbed parameter, $\rho: \beta_C$<br>Perturbed size, $\rho': 1.05 \cdot \beta_C$<br>Evaluation time, $t_n: [1, 1100] \text{ min}$ | $\beta_1 = 1 \text{ min}^{-1}$, $S = 300 \text{ nM}$, $\eta = 0.01 \text{ nM}^{-1} \text{ min}^{-1}$,<br>$\beta_2 = 0.3 \text{ min}^{-1}$, $\beta_I = 0.06 \text{ min}^{-1}$, $\beta_P = 0 \text{ min}^{-1}$,<br>$\beta_D = 0 \text{ min}^{-1}$, $\beta_A = 1.5 \text{ min}^{-1}$, $\gamma_A = 1.5 \text{ min}^{-1}$,<br>$K_A = 1 \text{ nM}$, $\gamma_{A0} = 0.1 \text{ min}^{-1}$, $\beta_M = 0.4167 \text{ min}^{-1}$,<br>$\gamma_M = 1.5 \text{ min}^{-1}$, $K_M = 1 \text{ nM}$, $\gamma_1 = 0.1 \text{ min}^{-1}$, $\beta_C = 0.06 \text{ min}^{-1}$, $\gamma_C = 0.1 \text{ min}^{-1}$, $\gamma_D = 0 \text{ min}^{-1}$ |
| **Fig. 2B** | Model: Eqs. S5 - S7<br>Controlled variable, $Y: X_C$<br>Perturbed parameter, $\rho: \beta_C$<br>Perturbed size, $\rho': 1.05 \cdot \beta_C$<br>Characterization parameter, $\theta: \beta_C$<br>Evaluation time, $t_n: [1, 1100] \text{ min}$ | $\beta_1 = 1 \text{ min}^{-1}$, $S = 300 \text{ nM}$, $\eta = 0.01 \text{ nM}^{-1} \text{ min}^{-1}$,<br>$\beta_2 = 0.3 \text{ min}^{-1}$, $\beta_I = 0.06 \text{ min}^{-1}$, $\beta_P = 0 \text{ min}^{-1}$, $\beta_D = 0 \text{ min}^{-1}$,<br>$\beta_A = 1.5 \text{ min}^{-1}$, $\gamma_A = 1.5 \text{ min}^{-1}$, $K_A = 1 \text{ nM}$,<br>$\gamma_{A0} = 0.1 \text{ min}^{-1}$, $\beta_M = 0.4167 \text{ min}^{-1}$, $\gamma_M = 1.5 \text{ min}^{-1}$,<br>$K_M = 1 \text{ nM}$, $\gamma_1 = 0.1 \text{ min}^{-1}$, $\beta_C = [0.01, 0.11] \text{ min}^{-1}$,<br>$\gamma_C = 0.1 \text{ min}^{-1}$, $\gamma_D = 0 \text{ min}^{-1}$ |
| **Fig. 2C** | Model: Eqs. S5 - S7<br>Controlled variable, $Y: X_C$<br>Perturbed parameter, $\rho: \beta_C$<br>Perturbed size, $\rho': 1.05 \cdot \beta_C$<br>Characterization parameter, $\theta: \beta_C$<br>Evaluation time, $t_n: \{50, 500\} \text{ min}$ | $\beta_1 = 1 \text{ min}^{-1}$, $S = 300 \text{ nM}$, $\eta = 0.01 \text{ nM}^{-1} \text{ min}^{-1}$,<br>$\beta_2 = 0.3 \text{ min}^{-1}$, $\beta_I = 0.06 \text{ min}^{-1}$, $\beta_P = 0 \text{ min}^{-1}$, $\beta_D = 0 \text{ min}^{-1}$,<br>$\beta_A = 1.5 \text{ min}^{-1}$, $\gamma_A = 1.5 \text{ min}^{-1}$, $K_A = 1 \text{ nM}$,<br>$\gamma_{A0} = 0.1 \text{ min}^{-1}$, $\beta_M = 0.4167 \text{ min}^{-1}$, $\gamma_M = 1.5 \text{ min}^{-1}$,<br>$K_M = 1 \text{ nM}$, $\gamma_1 = 0.1 \text{ min}^{-1}$, $\beta_C = [0.01, 0.11] \text{ min}^{-1}$,<br>$\gamma_C = 0.1 \text{ min}^{-1}$, $\gamma_D = 0 \text{ min}^{-1}$ |
| **Fig. 3B-F** | Model: Eqs. S5 - S7<br>Controlled variable, $Y: X_C$<br>Perturbed parameter, $\rho: \beta_C$<br>Perturbed size, $\rho': 1.05 \cdot \beta_C$<br>Evaluation time, $t_n: [0.1, 8000] \text{ min}$ | $\beta_1 = 1 \text{ min}^{-1}$, $S = 300 \text{ nM}$, $\eta = 0.01 \text{ nM}^{-1} \text{ min}^{-1}$, $\beta_2 = 0.3 \text{ min}^{-1}$,<br>$\beta_I = 0.06 \text{ min}^{-1}$, $\beta_P = \{0, 0.5\} \text{ min}^{-1}$, $\beta_D = \{0, 0.185\} \text{ min}^{-1}$,<br>$\beta_A = 1.5 \text{ min}^{-1}$, $\gamma_A = 1.5 \text{ min}^{-1}$, $K_A = 1 \text{ nM}$, $\gamma_{A0} = 0.1 \text{ min}^{-1}$,<br>$\beta_M = 0.4167 \text{ min}^{-1}$, $\gamma_M = 1.5 \text{ min}^{-1}$, $K_M = 1 \text{ nM}$, $\gamma_1 = 0.1 \text{ min}^{-1}$,<br>$\beta_C = 0.1 \text{ min}^{-1}$, $\gamma_C = 0.1 \text{ min}^{-1}$, $\gamma_D = 0 \text{ min}^{-1}$ |
| **Fig. 4A-C** | Model: Eqs. S5 - S7<br>Controlled variable, $Y: X_C$<br>Perturbed parameter, $\rho: \beta_C$<br>Perturbed size, $\rho': 1.05 \cdot \beta_C$<br>Evaluation time, $t_n: [15, 3000] \text{ min}$ | $\beta_1 = 1 \text{ min}^{-1}$, $S = 300 \text{ nM}$, $\eta = 0.01 \text{ nM}^{-1} \text{ min}^{-1}$, $\beta_2 = 0.3 \text{ min}^{-1}$,<br>$\beta_I = [0.02, 0.1] \text{ min}^{-1}$, $\beta_P = [0, 0.5] \text{ min}^{-1}$, $\beta_D = [0, 0.8] \text{ min}^{-1}$,<br>$\beta_A = 1.5 \text{ min}^{-1}$, $\gamma_A = 1.5 \text{ min}^{-1}$, $K_A = 1 \text{ nM}$, $\gamma_{A0} = 0.1 \text{ min}^{-1}$,<br>$\beta_M = 0.4167 \text{ min}^{-1}$, $\gamma_M = 1.5 \text{ min}^{-1}$, $K_M = 1 \text{ nM}$, $\gamma_1 = 0.1 \text{ min}^{-1}$,<br>$\beta_C = 0.1 \text{ min}^{-1}$, $\gamma_C = 0.1 \text{ min}^{-1}$, $\gamma_D = 0 \text{ min}^{-1}$ |
| **Fig. 5A-E** | Model: Eqs. S5 - S7<br>Controlled variable, $Y: X_C$<br>Perturbed parameter, $\rho: \beta_C$<br>Perturbed size, $\rho': 1.05 \cdot \beta_C$<br>Evaluation time, $t_n: [1, 5000] \text{ min}$ | $\beta_1 = 1 \text{ min}^{-1}$, $S = 300 \text{ nM}$, $\eta = 0.01 \text{ nM}^{-1} \text{ min}^{-1}$, $\beta_2 = 0.3 \text{ min}^{-1}$,<br>$\beta_I = 0.06 \text{ min}^{-1}$, $\beta_P = 0.25 \text{ min}^{-1}$, $\beta_D = 0.4 \text{ min}^{-1}$,<br>$\beta_A = 1.5 \text{ min}^{-1}$, $\gamma_A = 1.5 \text{ min}^{-1}$, $K_A = 1 \text{ nM}$, $\gamma_{A0} = 0.1 \text{ min}^{-1}$,<br>$\beta_M = 0.4167 \text{ min}^{-1}$, $\gamma_M = 1.5 \text{ min}^{-1}$, $K_M = 1 \text{ nM}$, $\gamma_1 = 0.1 \text{ min}^{-1}$,<br>$\beta_C = 0.1 \text{ min}^{-1}$, $\gamma_C = 0.1 \text{ min}^{-1}$, $\gamma_D = 0 \text{ min}^{-1}$ |

**Table S3.** Used parameter values in supplementary figures.

| Figure | CoRaDyn parameters | Model parameters |
| --- | --- | --- |
| **Fig. S1** | Model: Eqs. S5 - S7<br>Controlled variable, $Y: X_C$<br>Perturbed parameter, $\rho: \beta_C$<br>Perturbed size, $\rho': [1.01, 1.05] \cdot \beta_C$ | $\beta_1 = 1 \text{ min}^{-1}$, $S = 300 \text{ nM}$, $\eta = 0.01 \text{ nM}^{-1} \text{ min}^{-1}$,<br>$\beta_2 = 0.3 \text{ min}^{-1}$, $\beta_I = 0.06 \text{ min}^{-1}$, $\beta_P = 0 \text{ min}^{-1}$,<br>$\beta_D = 0 \text{ min}^{-1}$, $\beta_A = 1.5 \text{ min}^{-1}$, $\gamma_A = 1.5 \text{ min}^{-1}$,<br>$K_A = 1 \text{ nM}$, $\gamma_{A0} = 0.1 \text{ min}^{-1}$, $\beta_M = 0.4167 \text{ min}^{-1}$,<br>$\gamma_M = 1.5 \text{ min}^{-1}$, $K_M = 1 \text{ nM}$, $\gamma_1 = 0.1 \text{ min}^{-1}$, $\beta_C = 0.1 \text{ min}^{-1}$, $\gamma_C = 0.1 \text{ min}^{-1}$, $\gamma_D = [0, 0.05] \text{ min}^{-1}$ |
| **Fig. S2** | Model: Eqs. S5 - S7<br>Controlled variable, $Y: X_C$<br>Perturbed parameter, $\rho: \beta_C$<br>Perturbed size, $\rho': 1.05 \cdot \beta_C$<br>Evaluation time, $t_n: 500 \text{ min}$ | $\beta_1 = 1 \text{ min}^{-1}$, $S = 300 \text{ nM}$, $\eta = 0.01 \text{ nM}^{-1} \text{ min}^{-1}$, $\beta_2 = 0.3 \text{ min}^{-1}$, $\beta_I = [0.02, 0.1] \text{ min}^{-1}$, $\beta_P = [0, 0.5] \text{ min}^{-1}$,<br>$\beta_D = [0, 0.8] \text{ min}^{-1}$, $\beta_A = 1.5 \text{ min}^{-1}$, $\gamma_A = 1.5 \text{ min}^{-1}$,<br>$K_A = 1 \text{ nM}$, $\gamma_{A0} = 0.1 \text{ min}^{-1}$, $\beta_M = 0.4167 \text{ min}^{-1}$,<br>$\gamma_M = 1.5 \text{ min}^{-1}$, $K_M = 1 \text{ nM}$, $\gamma_1 = 0.1 \text{ min}^{-1}$, $\beta_C = 0.1 \text{ min}^{-1}$, $\gamma_C = 0.1 \text{ min}^{-1}$, $\gamma_D = 0 \text{ min}^{-1}$ |
| **Fig. S3 - S15** | Model: Eqs. S5 - S7<br>Controlled variable, $Y: X_C$<br>Perturbed parameter, $\rho: \beta_C$<br>Perturbed size, $\rho': 1.05 \cdot \beta_C$<br>Evaluation time, $t_n: [15, 3000] \text{ min}$ | $\beta_1 = 1 \text{ min}^{-1}$, $S = 300 \text{ nM}$, $\eta = 0.01 \text{ nM}^{-1} \text{ min}^{-1}$, $\beta_2 = 0.3 \text{ min}^{-1}$, $\beta_I = [0.02, 0.1] \text{ min}^{-1}$, $\beta_P = [0, 0.5] \text{ min}^{-1}$,<br>$\beta_D = [0, 0.8] \text{ min}^{-1}$, $\beta_A = 1.5 \text{ min}^{-1}$, $\gamma_A = 1.5 \text{ min}^{-1}$,<br>$K_A = 1 \text{ nM}$, $\gamma_{A0} = 0.1 \text{ min}^{-1}$, $\beta_M = 0.4167 \text{ min}^{-1}$,<br>$\gamma_M = 1.5 \text{ min}^{-1}$, $K_M = 1 \text{ nM}$, $\gamma_1 = 0.1 \text{ min}^{-1}$, $\beta_C = 0.1 \text{ min}^{-1}$, $\gamma_C = 0.1 \text{ min}^{-1}$, $\gamma_D = 0 \text{ min}^{-1}$ |
| **Fig. S16A-E** | Model: Eqs. S5 - S7<br>Controlled variable, $Y: X_C$<br>Perturbed parameter, $\rho: \beta_C$<br>Perturbed size, $\rho': 1.05 \cdot \beta_C$<br>Evaluation time, $t_n: [1, 5000] \text{ min}$ | $\beta_1 = 1 \text{ min}^{-1}$, $S = 300 \text{ nM}$, $\eta = 0.01 \text{ nM}^{-1} \text{ min}^{-1}$,<br>$\beta_2 = 0.3 \text{ min}^{-1}$, $\beta_I = 0.1 \text{ min}^{-1}$, $\beta_P = 0.25 \text{ min}^{-1}$,<br>$\beta_D = 0.4 \text{ min}^{-1}$, $\beta_A = 1.5 \text{ min}^{-1}$, $\gamma_A = 1.5 \text{ min}^{-1}$,<br>$K_A = 1 \text{ nM}$, $\gamma_{A0} = 0.1 \text{ min}^{-1}$, $\beta_M = 0.4167 \text{ min}^{-1}$,<br>$\gamma_M = 1.5 \text{ min}^{-1}$, $K_M = 1 \text{ nM}$, $\gamma_1 = 0.1 \text{ min}^{-1}$, $\beta_C = 0.1 \text{ min}^{-1}$, $\gamma_C = 0.1 \text{ min}^{-1}$, $\gamma_D = 0 \text{ min}^{-1}$ |

## References

1. Muzzey D, Gómez-Uribe CA, Mettetal JT, Van Oudenaarden A. A Systems-Level Analysis of Perfect Adaptation in Yeast Osmoregulation. Cell. 2009;138(1):160–171. doi:10.1016/j.cell.2009.04.047.

2. Brownlee C, Wheeler GL. Cellular calcium homeostasis and regulation of its dynamic perturbation. Quantitative Plant Biology. 2025;6:e5. doi:10.1017/qpb.2025.2.

3. Casey JR, Grinstein S, Orlowski J. Sensors and regulators of intracellular pH. Nature Reviews Molecular Cell Biology. 2010;11(1):50–61. doi:10.1038/nrm2820.

4. El-Samad H. Biological feedback control—Respect the loops. Cell Systems. 2021;12(6):477–487. doi:10.1016/j.cels.2021.05.004.

5. Gómez-Schiavon M, El-Samad H. CoRa—A general approach for quantifying biological feedback control. Proceedings of the National Academy of Sciences. 2022;119(36):e2206825119. doi:10.1073/pnas.2206825119.

6. Chevalier M, Gómez-Schiavon M, Ng AH, El-Samad H. Design and Analysis of a Proportional-Integral-Derivative Controller with Biological Molecules. Cell Systems. 2019;9(4):338–353.e10. doi:10.1016/j.cels.2019.08.010.

7. Cannon WB. ORGANIZATION FOR PHYSIOLOGICAL HOMEOSTASIS. Physiological Reviews. 1929;9(3):399–431. doi:10.1152/physrev.1929.9.3.399.

8. Somvanshi PR, Patel AK, Bhartiya S, Venkatesh KV. Implementation of integral feedback control in biological systems. WIREs Systems Biology and Medicine. 2015;7(5):301–316. doi:10.1002/wsbm.1307.

9. Koshland DE, Goldbeter A, Stock JB. Amplification and Adaptation in Regulatory and Sensory Systems. Science. 1982;217(4556):220–225.

10. Ma W, Trusina A, El-Samad H, Lim WA, Tang C. Defining Network Topologies that Can Achieve Biochemical Adaptation. Cell. 2009;138(4):760–773. doi:10.1016/j.cell.2009.06.013.

11. Purvis J, Lahav G. Encoding and Decoding Cellular Information through Signaling Dynamics. Cell. 2013;152(5):945–956. doi:10.1016/j.cell.2013.02.005.

12. Nakakuki T, Birtwistle MR, Saeki Y, Yumoto N, Ide K, Nagashima T, et al. Ligand-Specific c-Fos Expression Emerges from the Spatiotemporal Control of ErbB Network Dynamics. Cell. 2010;141(5):884–896. doi:10.1016/j.cell.2010.03.054.

13. Jiang Y, AkhavanAghdam Z, Tsimring LS, Hao N. Coupled feedback loops control the stimulus-dependent dynamics of the yeast transcription factor Msn2. Journal of Biological Chemistry. 2017;292(30):12366–12372. doi:10.1074/jbc.C117.800896.

14. Purvis JE, Karhohs KW, Mock C, Batchelor E, Loewer A, Lahav G. p53 Dynamics Control Cell Fate. Science. 2012;336(6087):1440–1444. doi:10.1126/science.1218351.

15. Sourjik V, Berg HC. Receptor sensitivity in bacterial chemotaxis. Proceedings of the National Academy of Sciences. 2002;99(1):123–127. doi:10.1073/pnas.011589998.

16. El-Samad H, Kurata H, Doyle JC, Gross CA, Khammash M. Surviving heat shock: Control strategies for robustness and performance. Proceedings of the National Academy of Sciences. 2005;102(8):2736–2741. doi:10.1073/pnas.0403510102.

17. Alves R, Savageau MA. Extending the method of mathematically controlled comparison to include numerical comparisons. Bioinformatics. 2000;16(9):786–798. doi:10.1093/bioinformatics/16.9.786.

18. Briat C, Gupta A, Khammash M. Antithetic Integral Feedback Ensures Robust Perfect Adaptation in Noisy Biomolecular Networks. Cell Systems. 2016;2(1):15–26. doi:10.1016/j.cels.2016.01.004.

19. Bennett S. Development of the PID controller. IEEE Control Systems. 1993;13(6):58–62. doi:10.1109/37.248006.

20. Thorsen K, Agafonov O, Selstø CH, Jolma IW, Ni XY, Drengstig T, et al. Robust Concentration and Frequency Control in Oscillatory Homeostats. PLoS ONE. 2014;9(9):e107766. doi:10.1371/journal.pone.0107766.

